# Salinity adaptation in date palm (*Phoenix dactylifera*): evidence for selective sweeps at Salt Overly Sensitive (SOS) genes

**DOI:** 10.64898/2026.11.03.762174

**Authors:** Peter Muchina, Shameem Younuskunju, Ayeda A. Ahmed, Yasmin A. Mohamoud, Joel A. Malek

## Abstract

The date palm is a high salinity tolerant fruit tree grown mainly in the region spanning from North Africa to parts of South Asia. Climate change models of this region indicate the possibility of increasing salinity in the future. The Salt Overly Sensitive (SOS) pathway is crucial for salinity tolerance in plants, serving as a primary defense mechanism by facilitating sodium exclusion. To better understand date palms’ salt tolerance, we analyzed variation in SOS pathway genes in a set of 197 date palm genomes collected from across the range of cultivation. Here, we show evidence for selective sweeps in multiple SOS pathway genes. A single major *pdSOS1* haplotype is at very high frequency in Eastern and Western subpopulations, with a minor, likely ancestral haplotype still present in both geographical locations. Allele-specific gene expression analysis showed differences in salt-treated roots between swept and ancestral haplotypes. Interestingly, sequence-based analysis of the *pdSOS1* haplotype suggested a second, much earlier introgression event between *P. theophrasti* and date palm, though direction is unclear. This event happened before both the selective sweep and the previously reported recent introgression to date palm. We identified a MITE transposable element inserted into a *pdSOS1* intron on the swept haplotype, and a similar insertion was found in an intron of the *SOS2* paralog *pdCIPK8*. Both insertions are unique to date palm among *Phoenix* species. The *pdCIPK8* haplotype containing the MITE is also at high frequency in both populations. Further analysis showed evidence for a selective sweep spanning the *pdSOS3* region in the western population that shares similarity to an Eastern haplotype spanning the promoter region only. Slower extended haplotype homozygosity decay surrounding the genes in the western population suggests that the swept haplotype originated in the East and spread to the West in all cases. Altogether, our data provide evidence of selective sweeps in the date palm SOS pathway and may go to explaining how this species has developed such high salinity tolerance.

## INTRODUCTION

Date palm (*Phoenix dactylifera L*.) is a major fruit crop of the arid and semi-arid regions stretching from North Africa to South Asia and is among the most salt-tolerant tree crops (Al Kharusi et al., 2017; Yaish & Kumar, 2015). Despite date palm’s significance and high salt tolerance, the mechanisms underlying this tolerance are poorly characterized. Different cultivars range significantly in their tolerance to salinity with the more tolerant varieties showing higher Na^+^ exclusion, higher K^+^/Na^+^ ratios, and better-preserved photosynthesis and water relations (Al Kharusi et al., 2017, 2019; Al-Dakheel & Hussain, 2026; Benaceur et al., 2024). Physiological and molecular studies show that date palm can tolerate seawater-level salinity through tissue-specific ion partitioning (Mueller et al., 2024; Oyarce et al., 2025; Pongrac et al., 2024) and growth in high salinity results in changes in both DNA methylation (Al-Harrasi et al., 2018) and the transcriptome, including alternative splicing (Xu et al., 2022; Yaish et al., 2017) and microRNAs (Yaish et al., 2015). These studies have typically only considered salinity response in a few region-specific cultivars and population-level variation that may have contributed to salt-adaptation in the species is unexplored.

Soil salinity and its negative effects on agricultural productivity is increasing as irrigation with brackish water and a changing climate degrade farmland, particularly in the arid and semi-arid regions where the date palm is cultivated (Ali et al., 2023; Zhang et al., 2022). Indeed, some modeling suggests that the Middle East could lose significant date palm production by 2070 due to climate change (Shabani et al., 2012). Because sodium becomes cytotoxic if it accumulates in the cell, a key part of plant salinity tolerance is keeping cytosolic Na^+^ low, either by limiting Na^+^ entry, transporting it to the vacuole, or actively exporting it out across the plasma membrane (Ji et al., 2013; Qiu et al., 2002). The Salt Overly Sensitive (SOS) pathway plays a central role in this export of sodium across the plasma membrane (Ali et al., 2023; Ji et al., 2013). It was first characterized in Arabidopsis and has 3 main component proteins that are highly conserved across many plant species. *SOS3*, a calcium-binding protein, senses the cytosolic Ca^2+^ signal caused by increased salt and interacts with *SOS2* to activate it. Activated *SOS2 (CIPK24)*, a serine/threonine protein kinase of the CBL-interacting protein kinase (CIPK) sub-family then phosphorylates *SOS1* (Ali et al., 2023). *SOS1* is a plasma-membrane Na^+^/H^+^ antiporter that is ultimately responsible for transporting the Na^+^ out of the cell (Qiu et al., 2002). There are other CIPK proteins that can activate SOS1 beyond SOS2 and in Arabidopsis *CIPK8* interacts with *CBL10* and *SOS1* to form a complex involved in salt response (Yin et al., 2020), and elevated *CIPK8* expression improves salt tolerance in other species (Ding et al., 2023). The conservation of SOS pathway genes and their involvement in sodium transport make them strong candidate genes for studying natural variation in salinity resistance. Previous studies have investigated the SOS pathway gene and protein expression in date palm salinity response (Pongrac et al., 2024; Xu et al., 2022) but none have examined potential selection in the population in this key pathway.

Date palm, a monocot, is a good candidate for understanding salinity resistance selection in cultivated plants: it was domesticated roughly 7,000 years ago in the Near East, is cultivated across a wide range, and shows clear genetic differentiation between eastern and western populations (Gros-Balthazard et al., 2018; Mathew et al., 2015; Mohamoud et al., 2019), partly from introgression with wild *Phoenix* relatives (Flowers et al., 2019). Previous studies showed potential selective sweeps or regions of selection in both eastern (36 loci) and western (20 loci) subpopulations with some indication of functional enrichment in those regions (Hazzouri et al., 2015) though none to our knowledge showed salinity-specific focus. A new, near complete reference date palm genome with full chromosome assembly of the Ajwa cultivar (Celii et al., 2026) provides an important starting point to better understand long-range evolutionary patterns in the date palm genome. Here we leverage genome-wide genotyping of almost 200 date samples that span cultivated date palms from Morocco to Pakistan, targeted long-read sequencing of specific cultivars, together with a dense panel of wild *Phoenix* species, to test whether *pdSOS1* and functionally linked genes bear signatures of selection. We report evidence for a selective sweep at *pdSOS1* with the candidate functional variation being a GC-rich, MITE-like insertion. We identify the same insertion within *pdCIPK8*, an *SOS2* paralog, where a possible soft sweep is detectable in the eastern date palm population. Lastly, we present evidence for a selective sweep that includes the *pdSOS3* gene region in the Western population with a related eastern haplotype in the gene regulatory region also elevated in frequency. Altogether these results provide evidence of selection in the SOS pathway as part of the history of salinity adaptation in date palm.

## MATERIALS AND METHODS

### Whole-genome sequencing data and variant calling

Whole-genome sequencing (WGS) datasets comprising 197 *Phoenix dactylifera* samples from our previous studies (Younuskunju et al., 2023, 2024) and 42 accessions representing other *Phoenix* species (Torres et al., 2018) were analyzed. The date palm dataset included commercially important cultivars collected across North Africa to South Asia (Mathew et al., 2015; Stephan et al., 2018). Raw reads were processed as described previously (Younuskunju et al., 2023) and aligned to the Ajwa reference genome PRJNA1207559;(Celii et al., 2026) using BWA-MEM2 (Li & Durbin, 2009). Duplicate reads were marked using the MarkDuplicatesSpark function in GATK v4.4.0.0 (McKenna et al., 2010). Variants were called for individual samples using DRAGEN v4.4.4 (Illumina, 2024) in gVCF mode, followed by joint genotyping to generate a combined VCF file. Raw SNPs were hard-filtered using bcftools v1.3.1 (Danecek et al., 2021) with the following parameters, QD < 2.0 || FS > 60.0 || MQ < 40.0 || SOR > 3.0 || MQRankSum < -12.5 || ReadPosRankSum < -8.0 || QUAL < 30 || INFO/DP > 17500 || INFO/DP < 1000 for downstream analysis.

### Long-read sequencing (ONT) and genome assembly

Genomic DNA was extracted from approximately 1 g of fresh leaf tissue using the Wizard HMW DNA Extraction Kit (Promega, A2920), with minor modifications to accommodate the increased leaf mass and preserve DNA integrity. To enrich long DNA fragments, fragments shorter than 25 kb were removed using the Short Fragment Eliminator Kit (EXP-SFE001, ONT) before library preparation. Sequencing libraries were prepared using the ligation sequencing kit V14 (SQK-LSK114, ONT) and loaded onto PromethION R10.4.1 flow cells (FLO-PRO114M, ONT). Raw FASTQ reads with a minimum quality score of Q10 and a read length greater than 500 bp were considered for genome assembly. Genome assembly was performed using Hifiasm v0.25.0 software, a haplotype resolved assembler (Cheng et al., 2021). Methylation status was called using Dorado v1.0.0 (Oxford Nanopore Technologies, 2025) with the all-context models matched to sequencing chemistry, --modified-bases-threshold 0 and re-basecalled from the POD5 over the region of interest. Reads with methylation information were assigned to haplotypes using cluster-specific SNPs and/or indels.

### Identification of SOS pathway homologues

Predicted protein sequences were extracted from the *Phoenix dactylifera* (Ajwa) genome annotation (Celii et al., 2026) using gffread (Pertea & Pertea, 2020). Arabidopsis thaliana SOS pathway proteins, including NHX7/*SOS1* (1,146 aa; UniProt accession Q9LKW9), *CIPK8* (445 aa; Q9STV4), and CBL4/*SOS3* (222 aa; O81223) were retrieved from UniProt (Bateman et al., 2017) and used as queries in BLASTP (Camacho et al., 2009) searches against the predicted date palm proteome, with an E-value threshold of 1 × 10^−10^. Candidate homologues were identified based on sequence similarity and alignment statistics, and the corresponding gene annotations were used to define their genomic coordinates and full gene spans. The identified candidates were designated *pdSOS1* (Chr07:32,649,282–32,697,765), *pdCIPK8* (Chr09:24,310,183–24,348,153) and *pdSOS3* (Chr09:28,769,234–28,775,762).

### Haplotype phasing and clustering

Following identification of the candidate SOS pathway genes, variants were extracted from chromosome 7 (*pdSOS1*), and chromosome 9 (*pdSOS3* and *pdCIPK8*). Date palm genotypes were phased with WhatsHap (Martin et al., 2023) and subsequently with Beagle (Browning et al., 2021). Phased haplotypes were visualized as heatmaps using the pheatmap package in R (Kolde, 2025). Clustering was performed on gene-specific regions: the coding sequence (CDS) of *pdSOS1*, a subregion of *pdCIPK8* (Chr09: 24,330,536–24,348,128), and a 4.5kb region upstream of *pdSOS3* transcription start site (TSS) (Chr09: 28,775,762–28,779,762). Pairwise Manhattan distances between phased haplotypes were used for hierarchical clustering with Ward’s minimum variance method (Ward.D2) (Strauss & von Maltitz, 2017), and clusters were defined from the resulting dendrograms. Representative haplotypes were identified by partitioning around medoids (PAM) (Van der Laan et al., 2003) and evaluated with pvclust using 1,000 multiscale bootstrap replicates, with cluster support reported as approximately unbiased (AU) and bootstrap probability (BP) values(Suzuki & Shimodaira, 2006).

Clustering was first performed within date palm to characterize haplotype diversity, and then repeated using SNPs fixed among other *Phoenix* species to resolve interspecific haplotype relationships, with particular attention to date palm haplotypes clustering with *P. theophrasti*. Haplotypes showing evidence of introgression were excluded from the selective-sweep analysis. For each individual, the cluster memberships of its two phased haplotypes were compared to characterize haplotype combinations.

### Selective-sweep analysis

Selective-sweep analyses were conducted separately for the *pdSOS1, pdCIPK8*, and *pdSOS3* regions using complementary population-genetic statistics implemented in Selscan v3.0.1 (Rahman et al., 2025a, 2025b; Szpiech, 2024; Szpiech & Hernandez, 2014), in each case excluding the *P. theophrasti*-associated date palm haplotypes identified by haplotype clustering. For *pdSOS1*, cross-population nSL (XP-nSL) (Szpiech et al., 2021) was computed between *P. dactylifera* and *P. sylvestris*. For *pdCIPK8*, within-population haplotype homozygosity was assessed using iHH12 (Garud et al., 2015) in *P. dactylifera*. For *pdSOS3*, cross-population EHH (XP-EHH) (Sabeti et al., 2007) was computed between Eastern and Western date palm populations.

Population differentiation (*F*_*ST*_) and nucleotide diversity (π) were estimated in 30-kb sliding windows with a 6-kb step size using VCFtools (Danecek et al., 2011). For *pdSOS1* and *pdCIPK8* these statistics were computed between *P. dactylifera* and *P. sylvestris*, whereas for *pdSOS3* they were computed between the Eastern and Western *P. dactylifera* populations, reflecting the intraspecific nature of the *pdSOS3* sweep. Fay and Wu’s H was computed using the calc_sfs_tests function in the R package rehh (Gautier et al., 2017) using P. roebelenii for ancestral-state polarization. Together, these statistics characterize patterns of genetic diversity and differentiation across the candidate-gene regions.

### Haplotype homozygosity decay

Extended haplotype homozygosity (EHH) (Sabeti et al., 2007) was computed around the focal SNP of each candidate region (*pdSOS1*, Chr07:32,679,064; *pdCIPK8*, Chr09:24,339,868; *pdSOS3*, Chr09:28,779,415, located in the *pdSOS3* upstream region) from the phased genotypes using selscan v1.2.0 (Szpiech & Hernandez, 2014) (--ehh, 100-kb window, cutoff 0.01), separately in the Eastern and Western date palm populations over an identical set of sites and after excluding *P. theophrasti*-associated haplotypes. EHH was tracked for the major allele, defined across the combined East–West cohort, so that both populations describe the same haplotype. To establish a neutral expectation, EHH was computed for 100 frequency-matched control SNPs from the same chromosome, each within ±3% of the focal major-allele frequency and excluding the centromere and a 250-kb interval around the focal site; background decay is summarized as the mean ± 95% confidence interval of these controls.

### Structural variation analysis

The genomic regions of *pdSOS1* (Chr07:32,649,282–32,697,765) and *pdCIPK8* (Chr09:24,310,183–24,348,153) were aligned to the long-read, haplotype-resolved genome assemblies described above using nucmer v3.1 (Kurtz et al., 2004) to identify the corresponding regions in each date palm cultivar. Multiple sequence alignments (MSAs) were generated using MAFFT (Katoh & Standley, 2013) of the identified regions and visually scanned for structural variants not identified in the VCF file. The analysis also included the current reference Barhee BC4 genome (PRJNA322046) and *P. roebelenii* (PRJNA629103), retrieved from NCBI. For *pdSOS3*, the repetitive element was searched against the Dfam 4.0 HMM database for date palm-specific repeats (Storer et al., 2021).

Haplotype-group assignments for the assembled genomes were determined using the SNP dataset described above. Each haplotype-resolved assembly was aligned to the reference genome using minimap2 v2.26 (Li, 2018), and genotypes at the haplotype-clustering SNP positions within the target regions were extracted from the resulting BAM alignments. A custom Perl script implemented an individual-versus-all comparison, comparing each ONT assembled haplotype independently with every haplotype clustered sample. These pairwise comparisons were used to determine the haplotype-group assignment of each ONT assembled haplotype.

### Functional analysis

The GC-rich insertion identified in the *pdSOS1* gene region was used to search the rest of the genome for other copies using BLASTN (E < 1e-5, single HSP ≥400 bp in length and similarity >70%). Gene expression data from the study of date palm under salinity conditions (SRA:ERP132137, EBI:E-MTAB-11001) was downloaded from the SRA and aligned to the Ajwa reference genome using STAAR 2.7.10b (Dobin et al., 2013). Counts per million (CPM) were documented for sense strand as the library was directional (Mueller et al., 2024). Reads could only be reliably assigned to haplotypes in the introns where SNP variation between haplotypes was high enough. Haplotype unique variants were used to assign reads to either the ancestral or swept haplotype based on consensus information from the haplotype clustering (see above). We used the ViennaRNA package (Lorenz et al., 2011) to fold the DNA surrounding the MITE insertion to calculate potential RNA structure minimum free energy at 37C compared to nucleotide composition controls of that region. Small RNA analysis was conducted on previously generated sequences from leaves and roots of Khalas seedlings grown in normal and salt conditions (Yaish et al., 2015) (SRA: SRP061704) Illumina small RNA adapters were removed and 18-30bp aligned to the Ajwa reference using Bowtie2.

## RESULTS

The analysis included short-read genome sequences from 239 *Phoenix* accessions, comprising 197 *P. dactylifera* individuals and 42 accessions representing other *Phoenix* species. We focused analysis on Chromosomes 7 and 9 which contain the SOS genes of interest. After filtering we obtained 1,876,172 and 1,981,898 SNPs for each chromosome respectively. The *pdSOS1, pdCIPK8* and *pdSOS3* analysis regions contained 4,619, 2,984 and 641 SNPs, respectively.

### Evidence of selection at *pdSOS1*

Hierarchical clustering of the phased *pdSOS1* coding sequences resolved four date palm haplotype groups among the 394 date palm haplotypes (Figure 1a; Supplementary Table 1). The *pdSOS1*_hap1.0 group was dominant, comprising 89.3% (352/394) of haplotypes and occurring across all sampled groups, Eastern (43.2%), Western (36.5%), Nile (10.5%) and introduced U.S. (9.9%) populations. The *pdSOS1*_hap2.1 group accounted for 5.6% (22/394) and was largely restricted to the Western population (86.4%), the remainder occurring in the introduced population. Introduced populations in the USA contain a mixture of Western and Eastern cultivars but might not have clear indication of origin. The *pdSOS1*_hap2.2 was rare (1.0%, 4/394) and confined to the Western population. A fourth, low-frequency group (3.8%, 15/394), interpreted the ancestral date palm haplotype (discussed below), was distributed across the Eastern (73.3%), Western (20.0%), and Nile (6.7%) populations.

**Figure 1.**
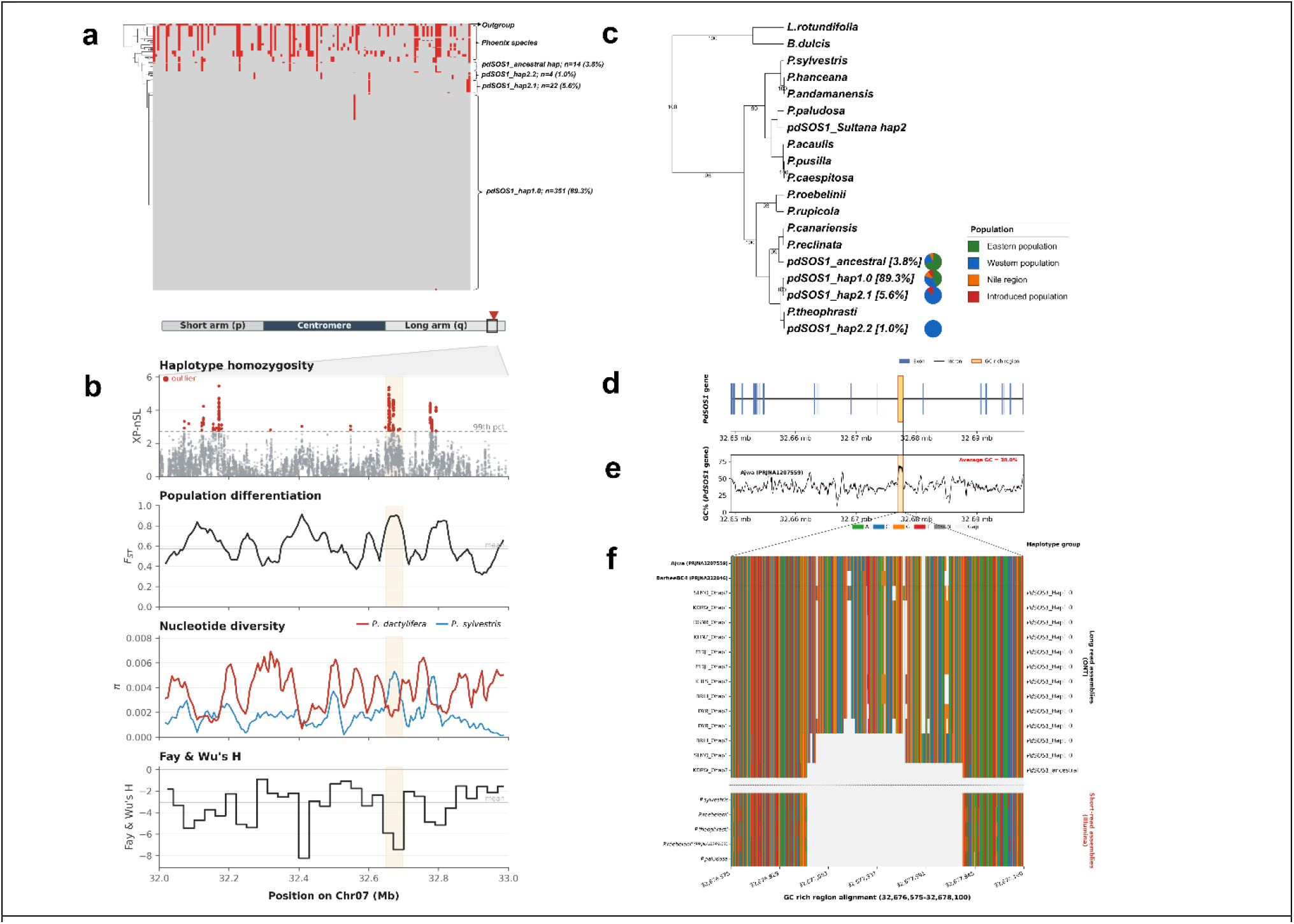
Haplotype structure and selection signatures at *pdSOS1*. (a) Heatmap of coding-sequence SNPs across *Phoenix* species and date palm haplotype clusters (individual haplotypes arranged horizontally; clustering dendrogram at left), revealing a single major haplotype present in approximately 90% of the date palm population. Grey indicates alleles matching the reference, while red indicates mismatches. (b) Selection statistics across Chr07:32–33 Mb; the shaded band marks the *pdSOS1* interval. From top: cross-population XP-nSL (*P. dactylifera* vs *P. sylvestris*; dashed line, 99th-percentile threshold = 2.73), fixation index (*F*_*ST*_), nucleotide diversity (π; *P. dactylifera* and *P. sylvestris*) and Fay & Wu’s *H*. (c) Haplotype dendrogram showing relationships among the four date palm *pdSOS1* haplotype groups, a divergent Sultana haplotype and other *Phoenix* species, with *L. rotundifolia* and *B. dulcis* included as a reference lineage; node values are bootstrap support (%). Pie charts indicate the population composition of the major haplotype groups. (d) Gene model of *pdSOS1* showing the position of the MITE (miniature inverted-repeat transposable element) insertion within an intron. (e) GC content across *pdSOS1* (average 38%), highlighting the GC-rich MITE insertion. (f) Alignment of the MITE insertion region across long-read (ONT) date palm assemblies and short-read (Illumina) *Phoenix* assemblies, showing sequence variation among swept (*pdSOS1*_hap1.0) haplotypes and absence of the insertion from the ancestral haplotype and other *Phoenix* species.

The haplotype heatmap revealed contrasting patterns of variation between the *P. dactylifera* haplotypes and other *Phoenix* species (Figure 1a). The relative homogeneity of *pdSOS1*_hap1.0 is consistent with reduced nucleotide diversity and the elevated XP-nSL signal observed at the *pdSOS1* locus. Indeed, the selection scan identified a localized signal overlapping *pdSOS1* on chromosome 7 (Chr07:32–33 Mb; Figure 1b). Because of the apparent hard sweep we used cross-population XP-nSL analysis comparing *P. dactylifera* with *P. sylvestris* which identified several sites exceeding the 99th-percentile threshold of 2.73, including sites within the *pdSOS1* region (Figure 1b). Genetic differentiation between the two species was elevated across chromosome 7, with a chromosome-wide mean *F*_*ST*_ of 0.572 and a local peak of 0.906 within *pdSOS1* (Figure 1b). Nucleotide diversity was generally higher in *P. dactylifera* than in *P. sylvestris* but declined locally in *P. dactylifera* across the *pdSOS1* region (Figure 1b). Fay and Wu’s *H* was negative across the region, with a chromosome-wide mean of −3.046 and a pronounced negative drop overlapping *pdSOS1*, reaching approximately −7.5 (Figure 1b). Together, elevated XP-nSL and *F*_*ST*_, a local reduction in nucleotide diversity and negative Fay and Wu’s *H* are consistent with a selection signal at the *pdSOS1* locus and the observation in the heatmap of reduced diversity across the entire population of date palm.

The haplotype dendrogram based on the distance matrix of the major *pdSOS1* haplotype groups revealed distinct relationships to other *Phoenix* lineages (Figure 1c). The predominant *pdSOS1*_hap1.0 group occupied a separate branch from the putative ancestral haplotype, whereas *pdSOS1*_hap2.1 and *pdSOS1*_hap2.2 were positioned close to *P. theophrasti*. Notably, *P. theophrasti* occurred within the broader date palm haplotype cluster rather than grouping with the other sampled *Phoenix* species. *pdSOS1_*hap*2*.2 likely represents the well-documented *P. theophrasti* introgression to date palm (Flowers et al., 2019); however, the fact that *P. theophrasti* appears as a sub-clade of date palm suggests a more complex history for this locus specifically (discussed below). We identified a putative ancestral haplotype characterized by its low frequency across the population and a lower number of derived mutations relative to the major date palm clade, rendering it less diverged from *Phoenix* species. We also observed a single eastern date palm cultivar, Sultana, that contained one haplotype clustering with other species. This likely represents a common eastern cultivar showing hybridization with a *Phoenix* species from the eastern contact zone.

### Functional analysis of the derived *pdSOS1* haplotype

With the most common derived haplotype (*pdSOS1*_hap1.0) appearing to have swept to high frequency in both the eastern and western date palm populations, we searched it for derived functional variants relative to the putative ancestral haplotype. Comparison of the coding sequence (NCBI predicted protein XP008798100, 1,153 aa) revealed no nonsense mutations and three nonsynonymous SNPs: Chr07:32,651,147 C>G (H949D), Chr07:32,649,614 G>A (D1081N) and Chr07:32,649,500 G>A (V1119I). Polarisation against the outgroup showed that the swept haplotype carries the derived allele at D1081N and V1119I, retaining the ancestral allele at H949D. All three lie in the C-terminal region outside the conserved cNMP-binding domain (aa 759–839), and none suggests an obvious functional change.

To look for regulatory differences, we generated haplotype-resolved long-read assemblies for eight cultivars (Barhi, Deglet Noor, Khenizi, Khalas, Medjool, Khadrawy, Salma and Dayri) and aligned the full *pdSOS1* region across all independently assembled haplotypes (Figure 1f). Although single-nucleotide variation among the assemblies was minimal, structural comparison identified an ∼750 bp GC-rich insertion in the intron (minus strand direction) of *pdSOS1* (LOC103713094), at Chr07:32,676,998–32,677,766 (769 bp) in the Ajwa reference and NC_052397.1:2,908,296–2,909,038 (743 bp) in the NCBI BarheeBC4 reference (Figure 1d-f). The insertion has a GC content of 67% versus 38% genome-wide (Figure 1e); its high GC content and palindromic sequence likely explain why it typically failed to sequence in short-read libraries, whereas long reads resolved both the sequence and breakpoints accurately. The insertion was present in most assembled haplotypes with a shorter variant in Barhi_hap1 (BRHI_Ohap2), Salma_hap1 (SLMQ_Ohap1), and completely absent from Khadrawy_hap2 (KDRW_Ohap2) (Figure 1). These structural states matched the SNP-based swept (*pdSOS1_h*ap1.0) and ancestral haplotype-group assignments. Analysis of the inserted sequence and its insertion site showed it to likely be a Miniature Inverted-Repeat Transposable Element (MITE) (Figure 2a, discussed below).

**Figure 2.**
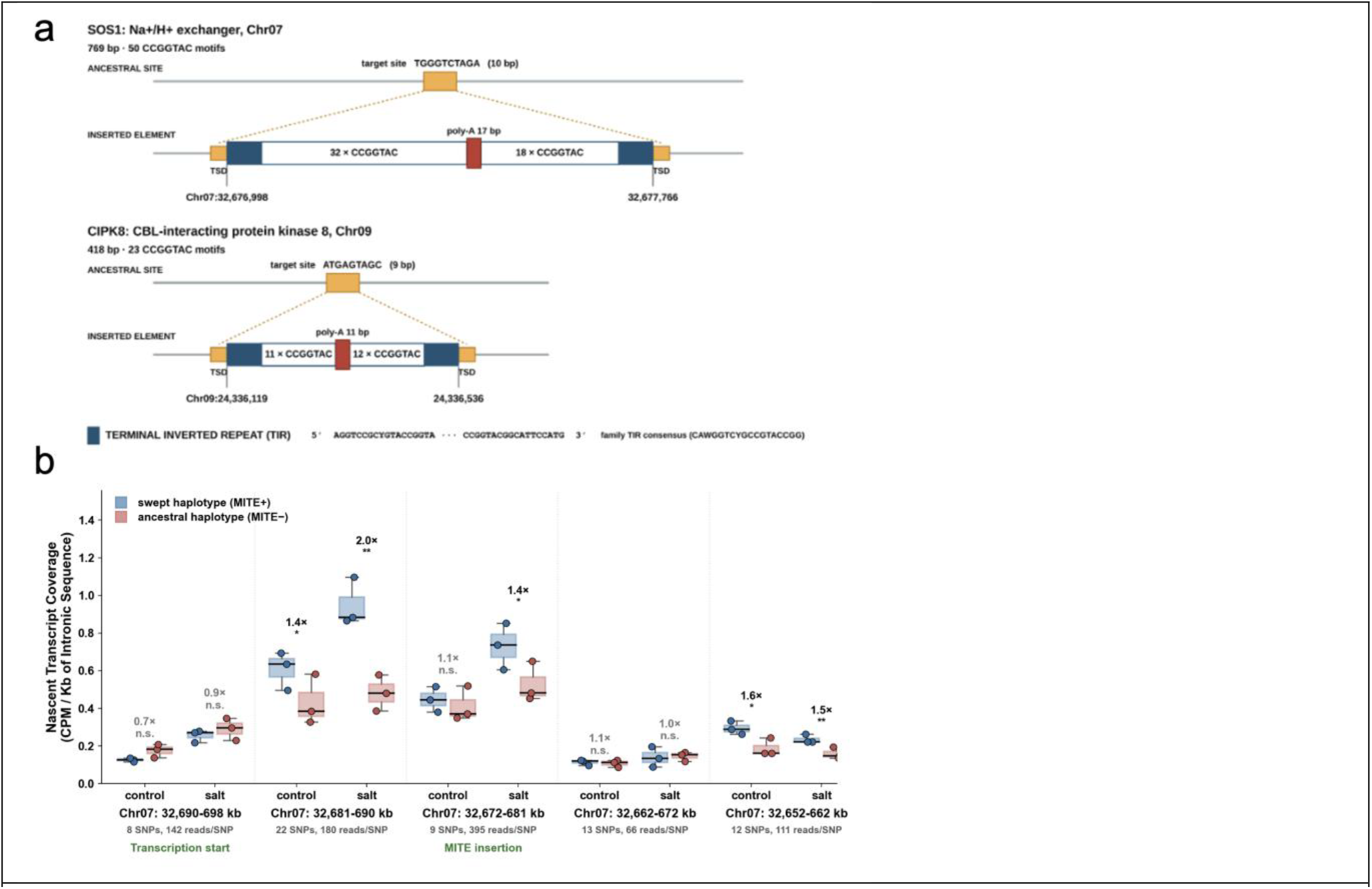
Expression and structure surrounding the MITE insertion in *pdSOS1* and *pdCIPK8*. (a) The MITE inserted in long introns of both *pdSOS1* and *pdCIPK8* and shows a similar structure including shared conserved Terminal Inverted Repeat sequences (TIR), a central poly-A stretch and varying levels of the repeating motif on either side. Other sequences are present in the two arms of the MITE, however the repeating motif of the CCGGTAC is the most prominent. *pdCIPK8’s* element ends exactly at CATTCCATG, while *pdSOS1* continues a further 12 bp (CGGCAGACCATG) before its target site duplication (TSD). The 3′ boundary is flush in *pdCIPK8* and extended in *pdSOS1*. The family TIR consensus was determined from the other instances identified by BLAST. (b) Gene expression data from date palm roots grown in control and high salinity conditions was analyzed for the cultivar Nabut Saif which contains both the ancestral (Mite-) and swept (Mite+) haplotypes. Allele-specific expression differences of nascent RNA (intronic) were observed with the swept haplotype demonstrating consistently higher levels of RNA mainly in two adjacent 8kb windows of the gene. Stars are significance based on paired T-test. Number of SNPs and aligned reads used to separate the two haplotypes is noted for each region. Note that genome coordinates are in reverse order to match the gene transcription direction.

To determine whether the inserted MITE arose on the date palm lineage, we assembled the corresponding region from the Illumina reads of the *Phoenix* species used for variant calling, including *P. reclinata, P. theophrasti, P. paludosa, P. sylvestris* and *P. roebelenii* (Methods) using SPAdes (Bankevich et al., 2012)(Supplementary Figure S1). All five species showed the uninserted configuration seen in the putative ancestral KDRW_hap2, and a direct search of the reads in the BAM files found no evidence of broken read pairs or clipped reads as seen in date palms with the MITE insertion (Figure 1f). Oxford Nanopore data showed no significant CG or CHH methylation differences between ancestral and derived haplotypes surrounding insertion (data not shown), though the DNA came from leaf tissue and may not reflect root methylation under salt stress.

To assess the effect of the swept haplotype on *pdSOS1* gene expression, we re-analysed total RNA-seq data from date palms grown under normal and high-salinity conditions (Mueller et al., 2024), which compared the salinity-tolerant Khalas with the intolerant Nabut Saif. Khalas is homozygous for the swept haplotype, and SNPs in the RNA-seq reads revealed that Nabut Saif is heterozygous for the swept and ancestral haplotypes, fortuitously allowing allele-specific analysis of the rare ancestral haplotype. Because haplotype-specific SNPs are rare in exons (Figure 1a), we measured nascent (intronic) mRNA in Nabut Saif. Both haplotypes were upregulated under salinity, and both appeared to initiate transcription at similar levels, but RNA levels diverged downstream, with the swept (MITE+) haplotype consistently showing significantly more RNA than the ancestral (MITE−) haplotype across multiple windows in roots (Figure 2b). Shoots showed the same trend at lower coverage and weaker significance (Supplementary Figure S2). Slower degradation or more stable polymerase elongation on the swept haplotype could account for this. Consistent with the latter, RNA folding predictions show the MITE insertion forms an extremely low minimum-free-energy structure (−563 kcal/mol), far below dinucleotide-shuffled controls (Z = −19.5; Supplementary Figure S3) and the resulting strong hairpins may alter RNA polymerase behavior and account for the higher RNA levels from this haplotype.

### Identification of selective sweeps in *pdCIPK8 and pdSOS3*

Following the identification of the MITE element in *pdSOS1*, we used BLAST to identify other locations in the genome containing the insertion. This resulted in 112 copies identified in 98 distinct genes (92% genic insertion), with *pdSOS1* being the longest hit among these. We searched the list of BLAST hits for other genes involved in the SOS pathway and identified *pdCIPK8* (a paralog of *SOS2*/CIPK24 (Yin et al., 2020)) in the group with a similarity of 74.8% to the MITE on a stretch of 453bp in *pdCIPK8’s* 9th intron. Comparing the *pdSOS1* and *pdCIPK8* insertions revealed a motif of CCGGTAC repeating multiple times interspersed among other sequences, including palindromes and a central Poly-A stretch. Further analysis showed evidence of a target duplication site (TSD) where the sequence inserted, suggesting it is likely a Miniature Inverted-repeat Transposable Element (MITE). The structure and terminal sequences of the putative MITE sequence were conserved between the *pdSOS1* and *pdCIPK8* sites, albeit of different lengths (Figure 2a).

With identification of the MITE in *pdCIPK8*, we wondered whether the shared event was random or an indication of possible coordinated selection. Hierarchical clustering of phased haplotypes at *pdCIPK8* again resolved a dominant group, *pdCIPK8*_hap1.0, present at high frequency (59.4%, 234/394) across both Eastern and Western populations (Figure 3a,c).

**Figure 3.**
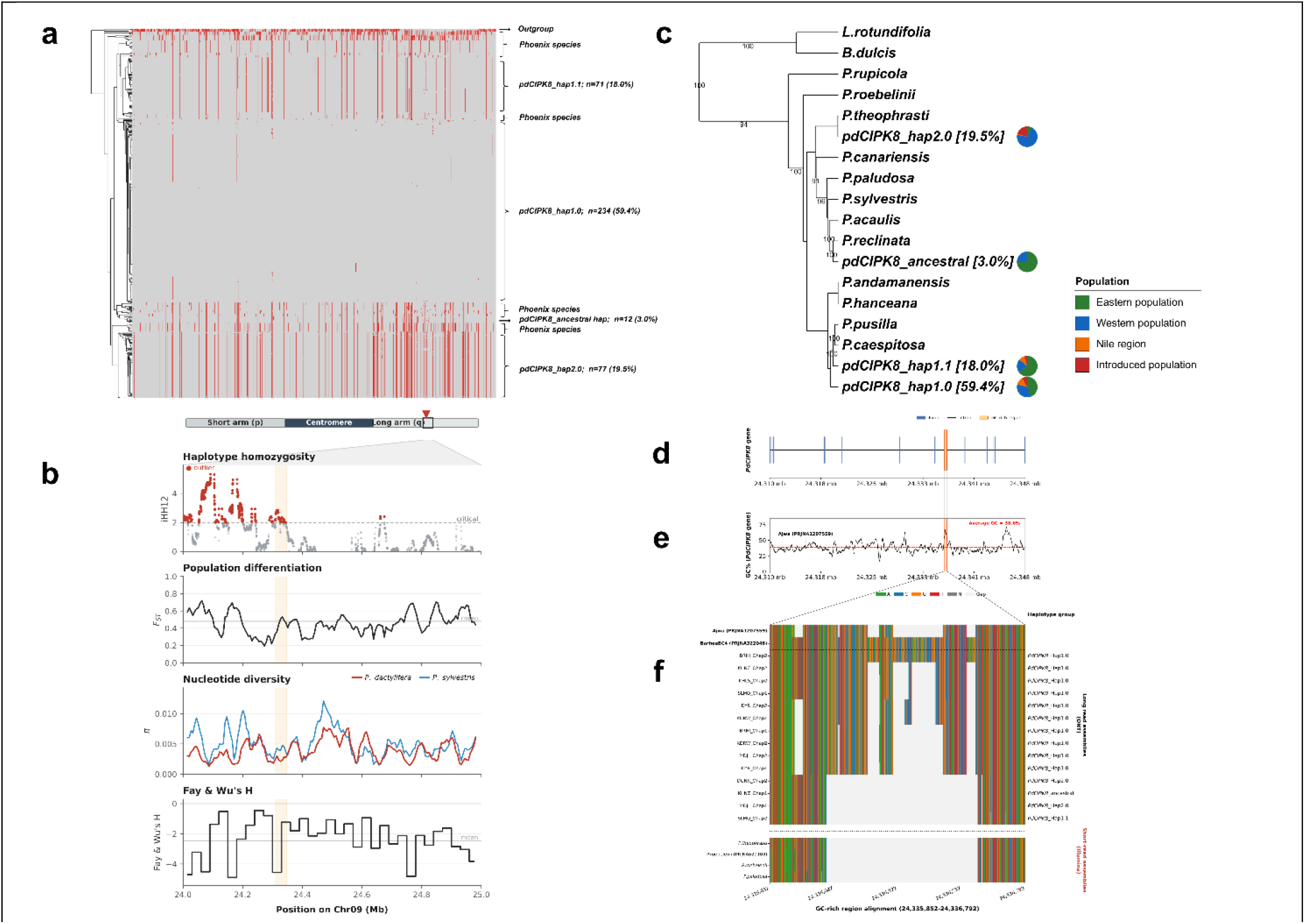
Haplotype structure and selection signatures at *pdCIPK8*. (a) Heatmap of SNPs across *pdCIPK8* haplotypes (individual haplotypes arranged horizontally; clustering dendrogram at left), showing the major date palm haplotype groups and other *Phoenix* species. Grey, alleles matching the reference; red, mismatches. (b) Selection statistics across Chr09:24–25 Mb; the shaded band marks the *pdCIPK8* interval. From top: within-population haplotype homozygosity (iHH12 in *P. dactylifera*; dashed line, significance threshold), fixation index (*F*_*ST*_), nucleotide diversity (π; *P. dactylifera* and *P. sylvestris*) and Fay & Wu’s *H*. (c) Haplotype dendrogram of the major date palm *pdCIPK8* haplotype groups and other *Phoenix* species, with *L. rotundifolia* and *B. dulcis* included as a reference lineage; node values are bootstrap support (%). Pie charts show the population composition of the major haplotype groups. (d) Gene model of *pdCIPK8* showing exons, introns and the position of the MITE (miniature inverted-repeat transposable element) insertion (e) GC content across *pdCIPK8* (average 38.6%), highlighting the GC-rich MITE insertion. (f) Alignment of the MITE insertion region across long-read (ONT) date palm assemblies and short-read (Illumina) *Phoenix* assemblies, showing presence of the insertion in the swept (*pdCIPK8*_hap1.0) haplotypes and its absence from the putative ancestral haplotype and other *Phoenix* species.

Within-population haplotype homozygosity (iHH12) was elevated across the locus (Figure 3b), the MITE-carrying haplotypes being the most common and occurring in both the Eastern and Western subpopulations; local *F*_*ST*_, reduced nucleotide diversity and negative Fay and Wu’s *H* were consistent with, though more modest than, the signal at *pdSOS1* (Figure 3b). Distance-based clustering identified a group likely introgressed from *P. theophrasti* (*pdCIPK8*_hap2.0) and placed the putative ancestral haplotype closer to other *Phoenix* species than the swept haplotypes (Figure 3c). As at *pdSOS1*, the derived swept haplotypes carried the MITE insertion, whereas the putative ancestral haplotype and other *Phoenix* species did not (Figure 3d–f).

The most common, putatively swept haplotype at *pdCIPK8* contained only one predicted non-synonymous SNP in the CDS at Chr09:24,342,437C>T resulting in R350C in the 411aa protein (XP038987688). That change sits within the CIPK_C domain which spans 277-389. However, during polarization of the SNP, we found the Cysteine to also be in *P. acaulis* as well as Brahea (*B. dulcis*) and Livistona (*L. rotundifolia*) outgroups with the remainder of *Phoenix* species containing the Arginine. Again, this left us with the MITE insertion as the best functional candidate to follow up on. Allele-specific gene expression was not possible for *pdCIPK8* because both date palm cultivars in the gene available expression data were homozygous for the swept haplotype. However, analysis of expression in both cultivars did show the expected trend of an increase in *pdCIPK8* expression in shoot but not root under salinity conditions (data not shown), agreeing with previous reports noting its role is more likely shoot focused (Yin et al., 2020).

With the discovery of putative selective sweeps at two SOS pathway related genes we searched the *SOS2* locus but did not find evidence of selection there. However, at the *pdSOS3* locus on Chr09 we detected evidence of selection specifically when comparing Eastern and Western date palm populations. Scans of the region using XP-EHH identified a long, high-frequency haplotype characteristic of a selective sweep in the Western population, supported by a local *F*_*ST*_ peak, reduced nucleotide diversity and more strongly negative Fay and Wu’s *H* in the West (Figure 4a). On inspection of the region in the Eastern population it became clear that the Western populations swept haplotype shared similarity to a specific Eastern haplotype that is also at higher frequency in the East but for which recombination had degraded it to containing mostly the promoter and upstream region of *pdSOS3* (Figure 4b).

**Figure 4.**
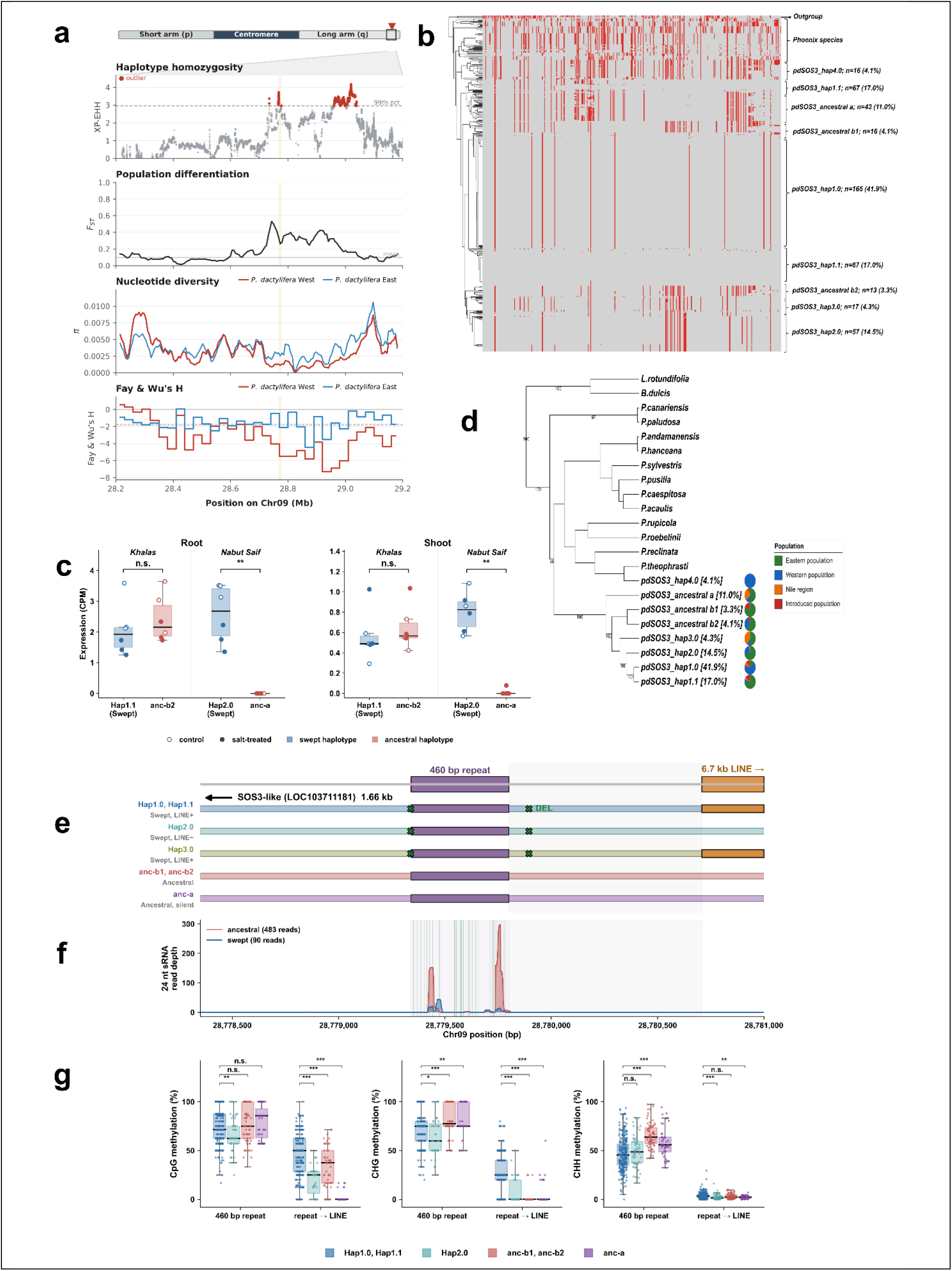
Analysis of the *pdSOS3* gene region. (a) Selection statistics across Chr09:28.2–29.2 Mb, all computed between Eastern and Western *P. dactylifera*; the shaded band marks the *pdSOS3* interval. From top: cross-population XP-EHH (dashed line, significance threshold), fixation index (*F*_*ST*_), nucleotide diversity (π) and Fay & Wu’s *H* (b) Heatmap of SNPs across *pdSOS3* haplotypes (individual haplotypes arranged horizontally; clustering dendrogram at left), showing the major date palm haplotype groups, with *Phoenix* species for comparison. Grey, alleles matching the reference; red, mismatches. (c) Gene expression analysis (SRA:ERP132137) of root and shoot in Khalas and Nabut Saif showed no detectable expression from the anc-a haplotype in any condition likely due to the SINE insertion near its TSS. (d) Haplotype dendrogram of the major date palm *pdSOS3* haplotype groups and other *Phoenix* species, with *L. rotundifolia* and *B. dulcis* included as a reference lineage; node values are bootstrap support (%). Pie charts show the population composition of the major haplotype groups. (e) The structure of the Dfam repeat (DR003745898.2) with 49 diagnostic snps separating the ancestral and swept haplotypes as green lines. (f) 24nt siRNAs from libraries of pooled Khalas root and shoot samples (SRA:SRP061704) under normal and salt conditions show alignment mainly to the ancestral haplotype in two regions of the repeat. The Dfam repeat is enriched for ancestral 24nt siRNAs vs surrounding 5kb region (70 SNP positions and only 6 reads aligned). All reads from 18-30bp were counted outside the Dfam region to detect haplotypes and a two-sided fisher exact test was used showing significantly more (p=5.3e-0.5) small RNA’s binding to the ancestral haplotype (483 reads) vs swept (90) in the Dfam repeat with 49 CHH sites of 460bp compared to outside element where only 6 swept haplotype reads aligned at 70 CHH sites within 5.2kb of sequence. Dashed line is the highest ancestral fraction possible given Khalas is known to be *pdSOS3*_hap1.1 and anc-b2. (g) CpG, CHG and CHH methylation on various haplotypes as determined by the Oxford Nanopore data. Note that the CHH methylation is higher in both ancestral haplotype groups specifically within the Dfam repeat which agrees with the 24nt sRNA alignment data.

The *pdSOS3* gene body was highly conserved across date palm haplotypes and *Phoenix* species with little variation between all haplotypes. However, the promoter region showed variation, including two deletions surrounding an interspersed repeat of 450bp (3200 copies in date palm, Dfam element DR003745898.2) at Chr09:28,779,351–28,779,801 and a LINE insertion approximately 3kb from the Transcription start of the *pdSOS3* gene. Analysis of the repeat in *Phoenix* species showed the first deletion is likely a target-site duplication of the repeat and variable across *Phoenix* species while the second deletion only occurs in date palm (Supplementary Figure S4). Interestingly, analysis with the plant genomic foundation model Botanic 1 (Barozet et al., 2026) showed the deleted region to be rich in plant regulatory elements (data not shown). Allele-specific gene expression analysis of the region showed mono-allelic expression from the swept haplotype when it was paired with an ancestral haplotype in Nabut Saif however that specific ancestral haplotype is one of 3 putative ancestral haplotypes and specifically contains a SINE insertion approximately 250bp from the transcription start (Figure 4c). Khalas contains a different ancestral haplotype (anc-b2) for which we did not detect differences in expression with the putatively swept haplotype (Figure 4c). Distance matrix analysis of the SNPs revealed a dendrogram with the expected introgressed *P. theophrasti* haplotype (*pdSOS3*_hap4.0) and a more complex clustering pattern than observed in *pdSOS1* and *pdCIPK8* however, the putative ancestral haplotype (anc-a) with the SINE insertion and no observed gene expression in the tested conditions was basal to the rest of the date palm haplotypes.

With no gene expression difference in the conditions tested between anc-b and the swept haplotype we searched the methylation results in the long-read data for possible differences among the various structural changes in the haplotypes (Figure 4e-g). We observed that the Dfam repeat element contained significantly less (two-sided Mann-Whitney U test) CHG and CHH methylation in the swept haplotypes (*pdSOS3*_hap1.0, hap1.1, hap2.0) vs the ancestral (anc-a, anc-b1, anc-b2), with the limitation that our long-read data did not have a representative of hap3.0 (Figure 4g). With the observed decrease in CHH methylation in the swept haplotypes we searched previous small RNA expression data (Yaish et al., 2015) for evidence of 24nt RNAs, a sign of possible RNA directed DNA methylation. Searching the region yielded significant 24nt RNA sequences aligning in two regions of the Dfam element in higher numbers on the ancestral haplotype than the swept haplotype (483 reads vs 90 reads on 49 haplotype distinguishing SNPs). Outside the region we loosened stringency to allow for any length of small RNA to determine the haplotypes contributing to the RNA (from pooled Khalas seedlings) and observed only 6 reads across 70 haplotype distinguishing SNPs and all 6 were swept haplotype derived. This is a statistically significant difference in ancestral to swept RNA production (Fisher exact p=5.3e-05) despite not knowing the exact frequency of each haplotype in the pool (Figure 4f) and may explain the higher CHH methylation in this region.

Comparison of extended haplotype homozygosity (EHH) decay around each swept focal SNP showed more rapid decay in the Eastern than the Western population at all three loci (Figure 5; ±100 kb shown in Supplementary Figure S5). Because the Western population has been shown to have longer background linkage disequilibrium, likely due to the *P. theophrasti* introgression (Flowers et al., 2019), introgressed haplotypes were excluded from all analyses; the elevated Western haplotype length at these loci therefore does not reflect introgression. For *pdSOS1* and *pdCIPK8*, the western and eastern haplotype homozygosity, and for the western in *pdSOS3*, decay was significantly longer than the allele-frequency matched chromosome background (Figure 5). The dissimilarity in the various gene location haplotype length in each population makes it more likely these were independent selection events rather than a universal elevated western haplotype length due to demography.

**Figure 5.**
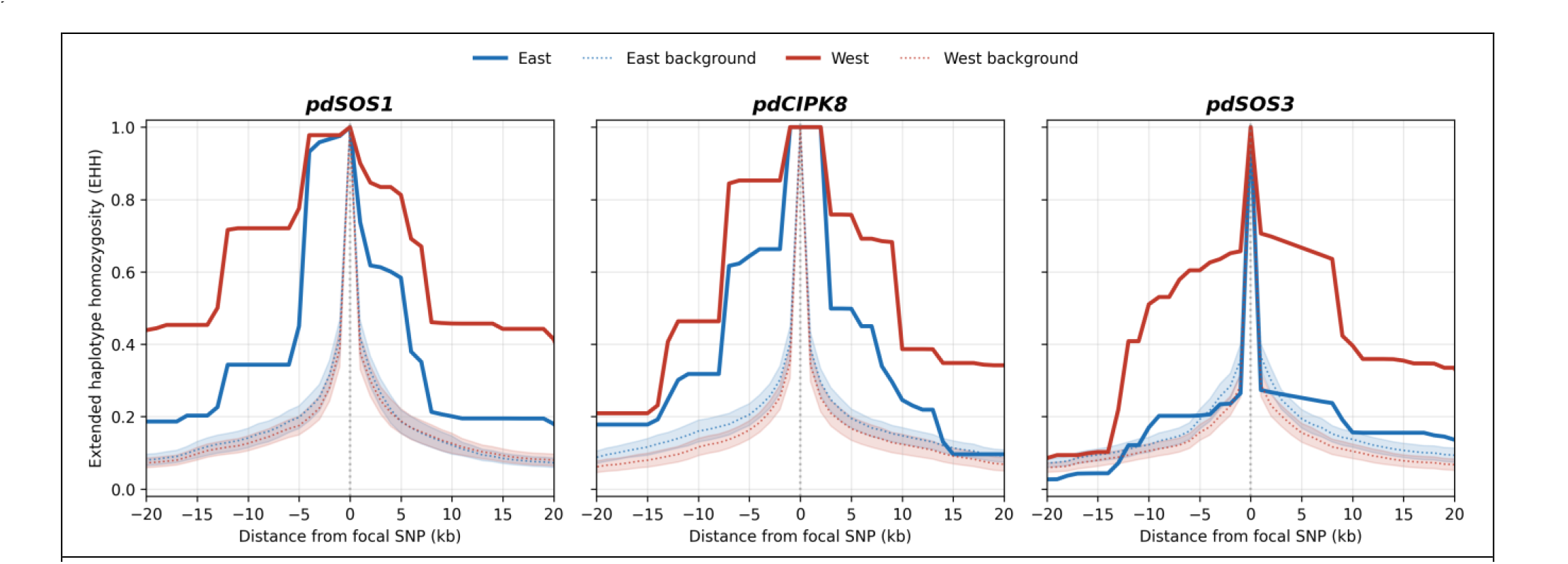
Extended haplotype homozygosity (EHH) for each gene region. Probability of homozygosity radiating from focal SNPs associated with the swept haplotype were plotted for each gene region; *pdSOS1* (Chr07:32,679,064), *pdCIPK8* (Chr09:24,339,868), and *pdSOS3* (Chr09:28,779,415). Solid lines, focal EHH; dotted lines and shaded bands, the mean and 95% confidence interval of 100 allele-frequency–matched control SNPs (within ±3% of the focal major-allele frequency) across the chromosome, excluding the centromere. In all populations except the Eastern at *pdSOS3*, the haplotypes are significantly longer than background and western haplotypes are longer than Eastern suggesting a spread from east to west. The varying eastern haplotype lengths suggest these arose from separate selection events rather than a single demographic event.

## DISCUSSION AND CONCLUSIONS

We present evidence for diverse selection patterns within the date palm SOS pathway, including both hard and soft sweeps across recent and ancient events. The results of these events may help explain the unique capacity of date palm to deal with high salinity conditions and could be critical to future breeding for improved salinity tolerance.

We began with the observation of a high-frequency, low-diversity haplotype spanning the long *pdSOS1* gene region. The extremely low diversity (Figure 1a) indicated this was likely a more recent event than the separation of the eastern and western subpopulations. The presence of an ancestral haplotype at low frequency in both subpopulations may either be balancing selection or an incomplete sweep. Interestingly, distance matrix analysis of the haplotypes revealed the ancestral date palm haplotype to be basal to a clade containing both *P. theophrasti* and its haplotype introgressed into date palm (Figure 1c). The structure of that relationship at this locus is unique, with *P. theophrasti* usually sitting outside the date palm clade for other loci (Figure 3c,4d) and previous studies (Torres et al., 2018). Though *P. theophrasti* does not have the MITE insertion of the date palm swept haplotype, the backbone is shared. For that reason, it is difficult to distinguish the direction of transfer of that haplotype. It could be either a much earlier event than the well-documented recent *P. theophrasti* introgression into date palm (Flowers et al., 2019) or it could have been the date palm introgressing *P. theophrasti*. Either one would have provided the backbone haplotype that the MITE inserted on. However, with the MITE-free haplotype at apparent high frequency in *P. theophrasti*, it would argue that some beneficial change had already risen on the backbone allowing it to increase in frequency regardless of the MITE insertion. Indeed, both haplotypes share the predicted derived D1044N amino acid change and further research would be needed to determine if this itself might be beneficial.

The discovery of a MITE insertion in *pdSOS1* led us to *pdCIPK8*, an *SOS2* paralog (Yin et al., 2020) where a related form inserted. It’s important to note that we used BLAST to search for the MITE sequence on the Ajwa reference (Celii et al., 2026) which could leave other instances undetected in other date palms, and a more complete population-wide search would be beneficial in the future. Both *pdSOS1* and *pdCIPK8* are involved in environmental stress response, which is interesting in light of previous findings of MITE involvement in stress response in Maize (Mao et al., 2015) and Apple (Niu et al., 2022). More importantly, a recent analysis of MITE insertions in multiple plant genomes showed that gene expression is higher for genes with MITE insertions, including insertions in the intron (Gao et al., 2025). Furthermore, a study of gene expression in rice under salinity stress identified a MITE insertion upstream of the *OsHKT1* potassium transporter and removal of the MITE reduced gene expression and increased salinity sensitivity (Wang et al., 2020). While the allele-specific expression analysis (Figure 2b) suggests that the MITE+ haplotype in *pdSOS1* has increased RNA-levels, because of the similarity between the two haplotypes in exonic regions, we cannot determine whether this higher level of nascent RNA translates to higher levels of spliced mRNA. The observation that nascent RNA is retained in two blocks near the MITE while being equal between the haplotypes at the beginning and end of the gene suggests a possible role of intron retention or RNA Pol II pausing specifically in the middle of the gene. Future work will be required to conduct long-read RNA-seq analysis under high-salt conditions to disambiguate the haplotype expression levels and their possible effect on protein expression. Our *pdSOS1* haplotype analysis of gene expression data showed concordance with a previous study of five cultivars, showing Khalas was salt tolerant while Nabut Saif was more susceptible (Al-Mulla et al., 2013) and for that reason they were selected by Mueller and colleagues for salinity studies (Maxim Messerer, 2023; Mueller et al., 2024). That Nabut Saif was selected for its salinity sensitivity and is heterozygous for the swept and ancestral *pdSOS1* haplotype is worth noting. Likewise, Pongrac et al 2024 and colleagues noted high gene expression differences between Sultana and Khalas both under control and salinity conditions and suggested they may be due to high genetic differences. This agrees with our findings that Sultana is likely a hybrid with another *Phoenix* species, with one of its haplotypes at *pdSOS1* being non-date palm. This may also explain why it demonstrated decreasing ability to avoid sodium translocation from roots to shoots with increasing irrigation salinity compared to the more salt-tolerant Khalas, which is homozygous for the swept *pdSOS1* haplotype (Pongrac et al., 2024) Whether the presence of the MITE in *pdSOS1* and *pdCIPK8* is simply a random event or represents a method for coordinated control between the genes requires further investigation. This control could happen through similar RNA structural change, as evidenced by the predicted very stable hairpin it forms (Supplementary Figure S3), or coordinated epigenetic modification of the region given the highly deviated nucleotide content.

For *pdSOS3*, we show clear evidence suggesting a selective sweep occurred in the Western population from a haplotype with Eastern origin that has since degraded in length by recombination. Whether the Eastern population had a much earlier sweep is not completely clear. The extended region of homozygosity in the Western haplotype includes the *pdSOS3* gene among others. Using the higher frequency but shorter haplotypes in the East identified the upstream 4.5kb (11kb including a 6.5kb inserted LINE) as the most likely region under selection in both populations. A deletion in that region is only present in swept haplotypes and their Eastern counterparts, while not present in other date palm haplotypes or *Phoenix* species. We did observe higher gene expression from the swept versus anc-a haplotype (Figure 4c), but that is a constitutive difference likely because anc-a contains a SINE insertion near the TSS. Whether the other anc-b has varying expression relative to the swept haplotype under certain conditions remains to be explored. We observed lower CHH and CHG methylation on the swept haplotypes in the Dfam element that borders the *Phoenix-specific* deletion, and this may play a role as a cis-regulator of *pdSOS3* gene expression. Lower CHH methylation tracked with fewer 24nt RNAs aligning in root expression analysis (Figure 4g). Indeed, lower CHH and CHG methylation could lead to more open chromatin and higher constitutive levels of *pdSOS3* expression in specific tissues and may provide a functional advantage of the swept haplotype in high salinity conditions.

Identifying genomic regions under selection will help in understanding date palm adaptation to its unique environment and could guide future breeding for climate change. Previous research identified 56 genomic regions that showed low nucleotide diversity in one date palm subpopulation or the other, providing evidence of region-specific selective sweeps (Hazzouri et al., 2015). However, in our findings, a recurring theme across the SOS pathway loci is the spread of putatively beneficial haplotypes from the eastern population into the western. In the case of *pdSOS1*, a small number of individuals in both populations retain ancestral haplotypes that are more closely related to each other than either is to the now dominant swept haplotype. They reveal that the two populations had likely already diverged prior to the sweep to near-fixation in both populations. This pattern could arise from the derived haplotype reaching the west by gene flow and then sweeping independently in each population, or from a single sweep in the east that subsequently spread westward. We propose the latter is more likely, as the swept haplotype appears essentially identical in both populations and a single origin fits the broader east-to-west pattern seen at other loci. In addition, we observed high diversity in the eastern population in the MITE insertions at both *pdSOS1* and *pdCIPK8*. Specifically, Iraqi varieties such as Barhi and Salma showed more divergence in their forms of the MITE with variable lengths and changes, and this high level of variation strengthens the likelihood that this region is the source of the original insertion. Either way, the derived *pdSOS1* haplotype was advantageous enough to displace the ancestral variation almost entirely in two already-diverged populations. This is an important consideration for future studies of regional selection in date palm, as these results suggest a pattern in which a beneficial allele arising in a distinct location (East) but with a similar climate challenge could spread across distant geographic regions, and this could happen at highly variable times in the population history. The methods to detect such adaptation would need to be adjusted depending on the history and geography of selection rather than any single method working universally.

Abiotic stress in plants is expected to increase in the future with climate change and date palm is unique in its capacity as a fruit tree to deal with high salinity. With this in mind, we investigated whether the classic SOS pathway in date palm showed evidence of selection. Our results show varying histories of apparent selection ranging from a hard sweep at *pdSOS1* to a possible soft sweep at *pdCIPK8* and a classic selective sweep signature only clear in the Western population for *pdSOS3* with the Eastern population showing evidence of the possibility of a much older sweep. We envision future studies testing whether plants homozygous for all swept haplotypes do indeed show significantly better salinity tolerance over those that do not. If this is the case, breeding for salinity adaptation could take advantage of this and further improve adaptation of this important tree.

## Data and code availability

- Phased haplotypes of the 197 date palm genomes at the *pdSOS1, pdCIPK8* and *pdSOS3* gene locations are available on request.
- Any additional information required to reanalyze the data reported in this paper is available from the lead contact upon request.

## Acknowledgments

Research funded by Qatar Research, Development and Innovation Council Grant (ARG02-0428-240366)

## Author contributions

Conceptualization, JAM and PM; sequencing and analysis SY, AAA, YAM; population analyses, PM, SY; writing, PM, JAM, SY, AAA.

## Declaration of generative AI and AI-assisted technologies

During the preparation of this work, the author(s) used Anthropic’s Claude to generate some of the figures from data, some literature review, and review the final text for suggested grammatical and syntax changes. After using this tool or service, the author(s) reviewed and edited the content as needed and take full responsibility for the content of the publication.

## REFERENCES

Al Kharusi, L., Assaha, D., Al-Yahyai, R., & Yaish, M. (2017). Screening of Date Palm (Phoenix dactylifera L.) Cultivars for Salinity Tolerance. Forests, 8(4), 136. 10.3390/f8040136

Al Kharusi, L., Sunkar, R., Al-Yahyai, R., & Yaish, M. W. (2019). Comparative Water Relations of Two Contrasting Date Palm Genotypes under Salinity. International Journal of Agronomy, 2019(1), 1–16. 10.1155/2019/4262013

Al-Dakheel, A. J., & Hussain, M. I. (2026). Differential physio-biochemical, mineral content, and fruit quality responses to water salinity in date palm (Phoenix dactylifera L.) varieties. Journal of Food Composition and Analysis, 152, 109024. 10.1016/j.jfca.2026.109024

Al-Harrasi, I., Al-Yahyai, R., & Yaish, M. W. (2018). Differential DNA methylation and transcription profiles in date palm roots exposed to salinity. PLOS ONE, 13(1), e0191492. 10.1371/journal.pone.0191492

Ali, A., Petrov, V., Yun, D.-J., & Gechev, T. (2023). Revisiting plant salt tolerance: novel components of the SOS pathway. Trends in Plant Science, 28(9), 1060–1069. 10.1016/j.tplants.2023.04.003

Al-Mulla, L., Bhat, N. R., & Khalil, M. (2013). Salt-tolerance of tissue-cultured date palm cultivars under controlled environment. Proceedings of World Academy of Science, Engineering and Technology, (80), 785.

Bankevich, A., Nurk, S., Antipov, D., Gurevich, A. A., Dvorkin, M., Kulikov, A. S., Lesin, V. M., Nikolenko, S. I., Pham, S., Prjibelski, A. D., Pyshkin, A. V, Sirotkin, A. V, Vyahhi, N., Tesler, G., Alekseyev, M. A., & Pevzner, P. A. (2012). SPAdes: a new genome assembly algorithm and its applications to single-cell sequencing. Journal of Computational Biology : A Journal of Computational Molecular Cell Biology, 19(5), 455–477. 10.1089/cmb.2012.0021

Barozet, A., Cabeli, V., Ogier du Terrail, J., Rukhovich, A., Janssoone, T., Klajer, G., Sheikhitarghi, Z., Andrews, G., Veran, C., & Strouk, L. (2026). BOTANIC-1: a series of long-context plant genomic foundation models in the agentic era. In bioRxiv (p. 2026.09.04.749355). Cold Spring Harbor Laboratory. 10.64898/2026.09.04.749355

Bateman, A., Martin, M. J., O’Donovan, C., Magrane, M., Alpi, E., Antunes, R., Bely, B., Bingley, M., Bonilla, C., Britto, R., Bursteinas, B., Bye-AJee, H., Cowley, A., Da Silva, A., De Giorgi, M., Dogan, T., Fazzini, F., Castro, L. G., Figueira, L., … Zhang, J. (2017). UniProt: the universal protein knowledgebase. Nucleic Acids Research, 45(D1), D158–D169. 10.1093/nar/gkw1099

Benaceur, I., Meziani, R., El Fadile, J., Hoinkis, J., Canas Kurz, E., Hellriegel, U., & Jaiti, F. (2024). Salt Stress Induces Contrasting Physiological and Biochemical Effects on Four Elite Date Palm Cultivars (Phoenix dactylifera L.) from Southeast Morocco. Plants, 13(2), 186. 10.3390/plants13020186

Browning, B. L., Tian, X., Zhou, Y., & Browning, S. R. (2021). Fast two-stage phasing of large-scale sequence data. The American Journal of Human Genetics, 108(10), 1880–1890. 10.1016/J.AJHG.2021.08.005

Camacho, C., Coulouris, G., Avagyan, V., Ma, N., Papadopoulos, J., Bealer, K., & Madden, T. L. (2009). BLAST+: architecture and applications. BMC Bioinformatics, 10(1), 421. 10.1186/1471-2105-10-421

Celii, M., Al-Bader, N., Mohammed, N., Zhou, Y., Fornasiero, A., Navarrete Rodriguez, M., Shuwaikan, R., Toor, U. A., Llaca, V., Fengler, K., Parakkal, P., Koo, D.-H., Gros-Balthazard, M., Battesti, V., Capote, T., Purugganan, M., Malek, J. A., Blilou, I., Poland, J., & Wing, R. A. (2026). Foundational genomic resources for date palm (Phoenix dactylifera): A gap-free, telomere-to-telomere phased assembly of Ajwa and 19 high-quality genome assemblies. Scientific Data 2026. 10.1038/S41597-026-08193-0

Cheng, H., Concepcion, G. T., Feng, X., Zhang, H., & Li, H. (2021). Haplotype-resolved de novo assembly using phased assembly graphs with hifiasm. Nature Methods, 18(2), 170–175. 10.1038/s41592-020-01056-5

Danecek, P., Auton, A., Abecasis, G., Albers, C. A., Banks, E., DePristo, M. A., Handsaker, R. E., Lunter, G., Marth, G. T., Sherry, S. T., McVean, G., & Durbin, R. (2011). The variant call format and VCFtools. Bioinformatics, 27(15), 2156–2158. 10.1093/BIOINFORMATICS/BTR330

Danecek, P., Bonfield, J. K., Liddle, J., Marshall, J., Ohan, V., Pollard, M. O., Whitwham, A., Keane, T., McCarthy, S. A., & Davies, R. M. (2021). Twelve years of SAMtools and BCFtools. GigaScience, 10(2), 1–4. 10.1093/GIGASCIENCE/GIAB008

Ding, X., Liu, B., Liu, H., Sun, X., Sun, X., Wang, W., & Zheng, C. (2023). A new CIPK gene CmCIPK8 enhances salt tolerance in transgenic chrysanthemum. Scientia Horticulturae, 308, 111562. 10.1016/j.scienta.2022.111562

Dobin, A., Davis, C. A., Schlesinger, F., Drenkow, J., Zaleski, C., Jha, S., Batut, P., Chaisson, M., & Gingeras, T. R. (2013). STAR: ultrafast universal RNA-seq aligner. Bioinformatics, 29(1), 15–21. 10.1093/bioinformatics/bts635

Flowers, J. M., Hazzouri, K. M., Gros-Balthazard, M., Mo, Z., Koutroumpa, K., Perrakis, A., Ferrand, S., Khierallah, H. S. M., Fuller, D. Q., Aberlenc, F., Fournaraki, C., & Purugganan, M. D. (2019). Cross-species hybridization and the origin of North African date palms. Proceedings of the National Academy of Sciences of the United States of America, 116(5), 1651–1658. 10.1073/PNAS.1817453116

Gao, J., Yang, L.-L., Wang, Y.-R., Zhao, Y.-Y., Shi, Y., Wei, S.-J., Chen, N., Zhang, Y.-L., Gao, W.-J., & Li, S.-F. (2025). MITE Annotation and Landscape in 207 Plant Genomes Reveal Their Evolutionary Dynamics and Functional Roles. Molecular Ecology Resources, 25(8), e70041. 10.1111/1755-0998.70041

Garud, N. R., Messer, P. W., Buzbas, E. O., & Petrov, D. A. (2015). Recent Selective Sweeps in North American Drosophila melanogaster Show Signatures of Soft Sweeps. PLOS Genetics, 11(2), e1005004. 10.1371/journal.pgen.1005004

Gautier, M., Klassmann, A., & Vitalis, R. (2017). rehh 2.0: a reimplementation of the R package rehh to detect positive selection from haplotype structure. Molecular Ecology Resources, 17(1), 78–90. 10.1111/1755-0998.12634

Gros-Balthazard, M., Hazzouri, K. M., & Flowers, J. M. (2018). Genomic Insights into Date Palm Origins. Genes, 9(10), 502. 10.3390/genes9100502

Hazzouri, K. M., Flowers, J. M., Visser, H. J., Khierallah, H. S. M., Rosas, U., Pham, G. M., Meyer, R. S., Johansen, C. K., Fresquez, Z. A., Masmoudi, K., Haider, N., El Kadri, N., Idaghdour, Y., Malek, J. A., Thirkhill, D., Markhand, G. S., Krueger, R. R., Zaid, A., & Purugganan, M. D. (2015). Whole genome re-sequencing of date palms yields insights into diversification of a fruit tree crop. Nature Communications, 6(1), 8824. 10.1038/ncomms9824

Illumina. (2024). DRAGEN secondary analysis | Software for NGS data analysis. https://www.illumina.com/products/by-type/informatics-products/dragen-secondary-analysis.html

Ji, H., Pardo, J. M., Batelli, G., Van Oosten, M. J., Bressan, R. A., & Li, X. (2013). The Salt Overly Sensitive (SOS) Pathway: Established and Emerging Roles. Molecular Plant, 6(2), 275–286. 10.1093/mp/sst017

Katoh, K., & Standley, D. M. (2013). MAFFT Multiple Sequence Alignment Software Version 7: Improvements in Performance and Usability. Molecular Biology and Evolution, 30(4), 772–780. 10.1093/molbev/mst010

Kolde, R. (2025). Pretty Heatmaps [R package pheatmap version 1.0.13]. CRAN: Contributed Packages. 10.32614/CRAN.PACKAGE.PHEATMAP

Kurtz, S., Phillippy, A., Delcher, A. L., Smoot, M., Shumway, M., Antonescu, C., & Salzberg, S. L. (2004). Versatile and open software for comparing large genomes. Genome Biology, 5(2), R12. 10.1186/gb-2004-5-2-r12

Li, H. (2018). Minimap2: pairwise alignment for nucleotide sequences. Bioinformatics, 34(18), 3094–3100. 10.1093/bioinformatics/bty191

Li, H., & Durbin, R. (2009). Fast and accurate short read alignment with Burrows–Wheeler transform. Bioinformatics, 25(14), 1754–1760. 10.1093/BIOINFORMATICS/BTP324

Lorenz, R., Bernhart, S. H., Höner Zu Siederdissen, C., Tafer, H., Flamm, C., Stadler, P. F., & Hofacker, I. L. (2011). ViennaRNA Package 2.0. Algorithms for Molecular Biology : AMB, 6(1), 26. 10.1186/1748-7188-6-26

Mao, H., Wang, H., Liu, S., Li, Z., Yang, X., Yan, J., Li, J., Tran, L.-S. P., & Qin, F. (2015). A transposable element in a NAC gene is associated with drought tolerance in maize seedlings. Nature Communications, 6(1), 8326. 10.1038/ncomms9326

Martin, M., Ebert, P., & Marschall, T. (2023). Read-Based Phasing and Analysis of Phased Variants with WhatsHap. Methods in Molecular Biology, 2590, 127–138. 10.1007/978-1-0716-2819-5_8/FIGURES/1

Mathew, L. S., Seidel, M. A., George, B., Mathew, S., Spannagl, M., Haberer, G., Torres, M. F., Al-Dous, E. K., Al-Azwani, E. K., Diboun, I., Krueger, R. R., Mayer, K. F. X., Mohamoud, Y. A., Suhre, K., & Malek, J. A. (2015). A genome-wide survey of date palm cultivars supports two major subpopulations in Phoenix dactylifera. G3: Genes, Genomes, Genetics, 5(7), 1429–1438. 10.1534/G3.115.018341/-/DC1

Maxim Messerer. (2023). RNAseq data of salt treated date palms (Phoenix dactylifera). BioStudies, E-MTAB-11001. https://www.ebi.ac.uk/biostudies/arrayexpress/studies/E-MTAB-11001

McKenna, A., Hanna, M., Banks, E., Sivachenko, A., Cibulskis, K., Kernytsky, A., Garimella, K., Altshuler, D., Gabriel, S., Daly, M., DePristo, M. A., McKenna, A., Hanna, M., Banks, E., Sivachenko, A., Cibulskis, K., Kernytsky, A., Garimella, K., Altshuler, D., … DePristo, M. A. (2010). The Genome Analysis Toolkit: A MapReduce framework for analyzing next-generation DNA sequencing data. Genome Research, 20(9), 1297–1303. 10.1101/GR.107524.110

Mohamoud, Y. A., Mathew, L. S., Torres, M. F., Younuskunju, S., Krueger, R., Suhre, K., & Malek, J. A. (2019). Novel subpopulations in date palm (Phoenix dactylifera) identified by population-wide organellar genome sequencing. BMC Genomics, 20(1), 498. 10.1186/s12864-019-5834-7

Mueller, H. M., Franzisky, B. L., Messerer, M., Du, B., Lux, T., White, P. J., Carpentier, S. C., Winkler, J. B., Schnitzler, J.-P., El-Serehy, H. A., Al-Rasheid, K. A. S., Al-Harbi, N., Alfarraj, S., Kudla, J., Kangasjärvi, J., Reichelt, M., Mithöfer, A., Mayer, K. F. X., Rennenberg, H., … Geilfus, C.-M. (2024). Integrative multi-omics analyses of date palm (Phoenix dactylifera) roots and leaves reveal how the halophyte land plant copes with sea water. The Plant Genome, 17(1), e20372. 10.1002/tpg2.20372

Niu, C., Jiang, L., Cao, F., Liu, C., Guo, J., Zhang, Z., Yue, Q., Hou, N., Liu, Z., Li, X., Tahir, M. M., He, J., Li, Z., Li, C., Ma, F., & Guan, Q. (2022). Methylation of a MITE insertion in the MdRFNR1-1 promoter is positively associated with its allelic expression in apple in response to drought stress. The Plant Cell, 34(10), 3983–4006. 10.1093/plcell/koac220 Oxford Nanopore Technologies. (2025). Dorado (Version 1.0.0) [Computer software]. https://github.com/nanoporetech/dorado

Oyarce, P., Xiao, T. T., Henkel, C., Frederiksen, S. F., Gonzalez-Kise, J. K., Smet, W., Wang, J. Y., Al-Babili, S., & Blilou, I. (2025). Microscopy and spatial-metabolomics identify tissue-specific metabolic pathways uncovering salinity and drought tolerance mechanisms in Avicennia marina and Phoenix dactylifera roots. Scientific Reports, 15(1), 1076. 10.1038/s41598-025-85416-1

Pertea, M., & Pertea, G. (2020). GFF Utilities: GffRead and GffCompare. F1000Research, 9, 304. 10.12688/F1000RESEARCH.23297.1

Pongrac, P., Fischer, S., Wright, G., Thompson, J. A., Kelemen, M., Neugebauer, K., Brown, L. K., Vogel-Mikuš, K., Šala, M., El-Serehy, H. A., & White, P. J. (2024). Element partitioning, element localisation and transcriptome responses of date palms exposed to NaCl. In bioRxiv (p. 2024.12.07.627310). Cold Spring Harbor Laboratory. 10.1101/2024.12.07.627310

Qiu, Q.-S., Guo, Y., Dietrich, M. A., Schumaker, K. S., & Zhu, J.-K. (2002). Regulation of SOS1, a plasma membrane Na+/H+ exchanger in Arabidopsis thaliana, by SOS2 and SOS3. Proceedings of the National Academy of Sciences of the United States of America, 99(12), 8436–8441. 10.1073/pnas.122224699

Rahman, A., Smith, T. Q., & Szpiech, Z. A. (2025a). Fast and Memory-Efficient Dynamic Programming Approach for Large-Scale EHH-Based Selection Scans. Molecular Biology and Evolution, 42(11). 10.1093/molbev/msaf275

Rahman, A., Smith, T. Q., & Szpiech, Z. A. (2025b). Selection scans and downstream analysis with selscan. BioRxiv, 2025.10.10.681670. 10.1101/2025.10.10.681670

Sabeti, P. C., Varilly, P., Fry, B., Lohmueller, J., Hostetter, E., Cotsapas, C., Xie, X., Byrne, E. H., McCarroll, S. A., Gaudet, R., Schaffner, S. F., & Lander, E. S. (2007). Genome-wide detection and characterization of positive selection in human populations. Nature, 449(7164), 913–918. 10.1038/nature06250

Shabani, F., Kumar, L., & Taylor, S. (2012). Climate change impacts on the future distribution of date palms: a modeling exercise using CLIMEX. PloS One, 7(10), e48021. 10.1371/journal.pone.0048021

Stephan, N., Halama, A., Mathew, S., Hayat, S., Bhagwat, A., Mathew, L. S., Diboun, I., Malek, J., & Suhre, K. (2018). A comprehensive metabolomic data set of date palm fruit. Data in Brief, 18, 1313–1321. 10.1016/J.DIB.2018.04.012

Storer, J., Hubley, R., Rosen, J., Wheeler, T. J., & Smit, A. F. (2021). The Dfam community resource of transposable element families, sequence models, and genome annotations. Mobile DNA, 12(1), 2. 10.1186/s13100-020-00230-y

Strauss, T., & von Maltitz, M. J. (2017). Generalising Ward’s Method for Use with Manhattan Distances. PLOS ONE, 12(1), e0168288. 10.1371/journal.pone.0168288

Suzuki, R., & Shimodaira, H. (2006). Pvclust: an R package for assessing the uncertainty in hierarchical clustering. Bioinformatics, 22(12), 1540–1542. 10.1093/bioinformatics/btl117

Szpiech, Z. A. (2024). selscan 2.0: scanning for sweeps in unphased data. Bioinformatics, 40(1). 10.1093/bioinformatics/btae006

Szpiech, Z. A., & Hernandez, R. D. (2014). selscan: An Efficient Multithreaded Program to Perform EHH-Based Scans for Positive Selection. Molecular Biology and Evolution, 31(10), 2824–2827. 10.1093/MOLBEV/MSU211

Szpiech, Z. A., Novak, T. E., Bailey, N. P., & Stevison, L. S. (2021). Application of a novel haplotype-based scan for local adaptation to study high-altitude adaptation in rhesus macaques. Evolution Letters, 5(4), 408–421. 10.1002/EVL3.232

Torres, M. F., Mathew, L. S., Ahmed, I., Al-Azwani, I. K., Krueger, R., Rivera-Nuñez, D., Mohamoud, Y. A., Clark, A. G., Suhre, K., & Malek, J. A. (2018). Genus-wide sequencing supports a two-locus model for sex-determination in Phoenix. Nature Communications, 9(1), 3969. 10.1038/s41467-018-06375-y

Van der Laan, M., Pollard, K., & Bryan, J. (2003). A new partitioning around medoids algorithm. Journal of Statistical Computation and Simulation, 73(8), 575–584. 10.1080/0094965031000136012

Wang, J., Nan, N., Li, N., Liu, Y., Wang, T.-J., Hwang, I., Liu, B., & Xu, Z.-Y. (2020). A DNA Methylation Reader-Chaperone Regulator-Transcription Factor Complex Activates OsHKT1;5 Expression during Salinity Stress. The Plant Cell, 32(11), 3535–3558. 10.1105/tpc.20.00301

Xu, Z., Zhang, N., Fu, H., Wang, F., Wen, M., Chang, H., Wu, J., Abdelaala, W. B., Luo, Q., Li, Y., Li, C., Wang, Q., & Wang, Z. Y. (2022). Salt Stress Modulates the Landscape of Transcriptome and Alternative Splicing in Date Palm (Phoenix dactylifera L.). Frontiers in Plant Science, 12, 807739. 10.3389/FPLS.2021.807739/TEXT

Yaish, M. W., & Kumar, P. P. (2015). Salt tolerance research in date palm tree (Phoenix dactylifera L.), past, present, and future perspectives. Frontiers in Plant Science, 6(MAY), 1–5. 10.3389/FPLS.2015.00348

Yaish, M. W., Patankar, H. V., Assaha, D. V. M., Zheng, Y., Al-Yahyai, R., & Sunkar, R. (2017). Genome-wide expression profiling in leaves and roots of date palm (Phoenix dactylifera L.) exposed to salinity. BMC Genomics, 18(1), 246. 10.1186/s12864-017-3633-6

Yaish, M. W., Sunkar, R., Zheng, Y., Ji, B., Al-Yahyai, R., & Farooq, S. A. (2015). A genome-wide identification of the miRNAome in response to salinity stress in date palm (Phoenix dactylifera L.). Frontiers in Plant Science, 6(NOVEMBER), 151180. 10.3389/FPLS.2015.00946/TEXT

Yin, X., Xia, Y., Xie, Q., Cao, Y., Wang, Z., Hao, G., Song, J., Zhou, Y., & Jiang, X. (2020). The protein kinase complex CBL10–CIPK8–SOS1 functions in Arabidopsis to regulate salt tolerance. Journal of Experimental Botany, 71(6), 1801–1814. 10.1093/jxb/erz549

Younuskunju, S., Mohamoud, Y. A., Mathew, L. S., Mayer, K. F. X., Suhre, K., & Malek, J. A. (2023). Genome-wide association of dry (Tamar) date palm fruit color. The Plant Genome, 16(4), e20373. 10.1002/tpg2.20373

Younuskunju, S., Mohamoud, Y. A., Mathew, L. S., Mayer, K. F. X., Suhre, K., & Malek, J. A. (2024). The genetics of fruit skin separation in date palm. BMC Plant Biology, 24(1), 1050. 10.1186/s12870-024-05713-1

Zhang, H., Zhu, J., Gong, Z., & Zhu, J.-K. (2022). Abiotic stress responses in plants. Nature Reviews Genetics, 23(2), 104–119. 10.1038/s41576-021-00413-0

