## Supplementary information for "Salinity adaptation in date palm (*Phoenix dactylifera*): evidence for selective sweeps at Salt Overly Sensitive (SOS) genes"

Supplementary Table 1. Geographic distribution of pdSOS1 CDS haplotypes across cultivated date palm populations.

| Population group | Country (ISO) | Hap 1.0 (*n*=352) | Hap 2.1 (*n*=22) | Hap 2.2 (*n*=4) | Ancestral (*n*=15) | Total |
|---|---|---|---|---|---|---|
| **Eastern population** | | **93.3%** *(43.2%)* | **0.0%** *(0.0%)* | **0.0%** *(0.0%)* | **6.7%** *(73.3%)* | 163 |
| | AE | 39 | 0 | 0 | 2 | 41 |
| | IQ | 22 | 0 | 0 | 0 | 22 |
| | IR | 7 | 0 | 0 | 1 | 8 |
| | JO | 9 | 0 | 0 | 1 | 10 |
| | OM | 14 | 0 | 0 | 0 | 14 |
| | PK | 9 | 0 | 0 | 1 | 10 |
| | QA | 12 | 0 | 0 | 2 | 14 |
| | SA | 40 | 0 | 0 | 4 | 44 |
| **Western population** | | **83.1%** *(36.5%)* | **12.3%** *(86.4%)* | **2.6%** *(100.0%)* | **1.9%** *(20.0%)* | 154 |
| | MA | 52 | 12 | 0 | 2 | 66 |
| | DZ | 57 | 4 | 3 | 0 | 64 |
| | TN | 11 | 1 | 1 | 1 | 14 |
| | LY | 8 | 2 | 0 | 0 | 10 |
| **Nile region** | | **97.4%** *(10.5%)* | **0.0%** *(0.0%)* | **0.0%** *(0.0%)* | **2.6%** *(6.7%)* | 38 |
| | EG | 23 | 0 | 0 | 1 | 24 |
| | SD | 14 | 0 | 0 | 0 | 14 |
| **Introduced population** | | **92.1%** *(9.9%)* | **7.9%** *(13.6%)* | **0.0%** *(0.0%)* | **0.0%** *(0.0%)* | 38 |
| | US | 35 | 3 | 0 | 0 | 38 |

*Note: Bold values in population-group rows represent the percentage distribution of haplotypes within each population group, with horizontal percentages summing to 100%. Values in italicized text indicate the contribution of each population group to the global pool of the corresponding haplotype, with vertical percentages summing to 100%. Country-level values represent the number of resolved haplotypes assigned to each haplotype group.*

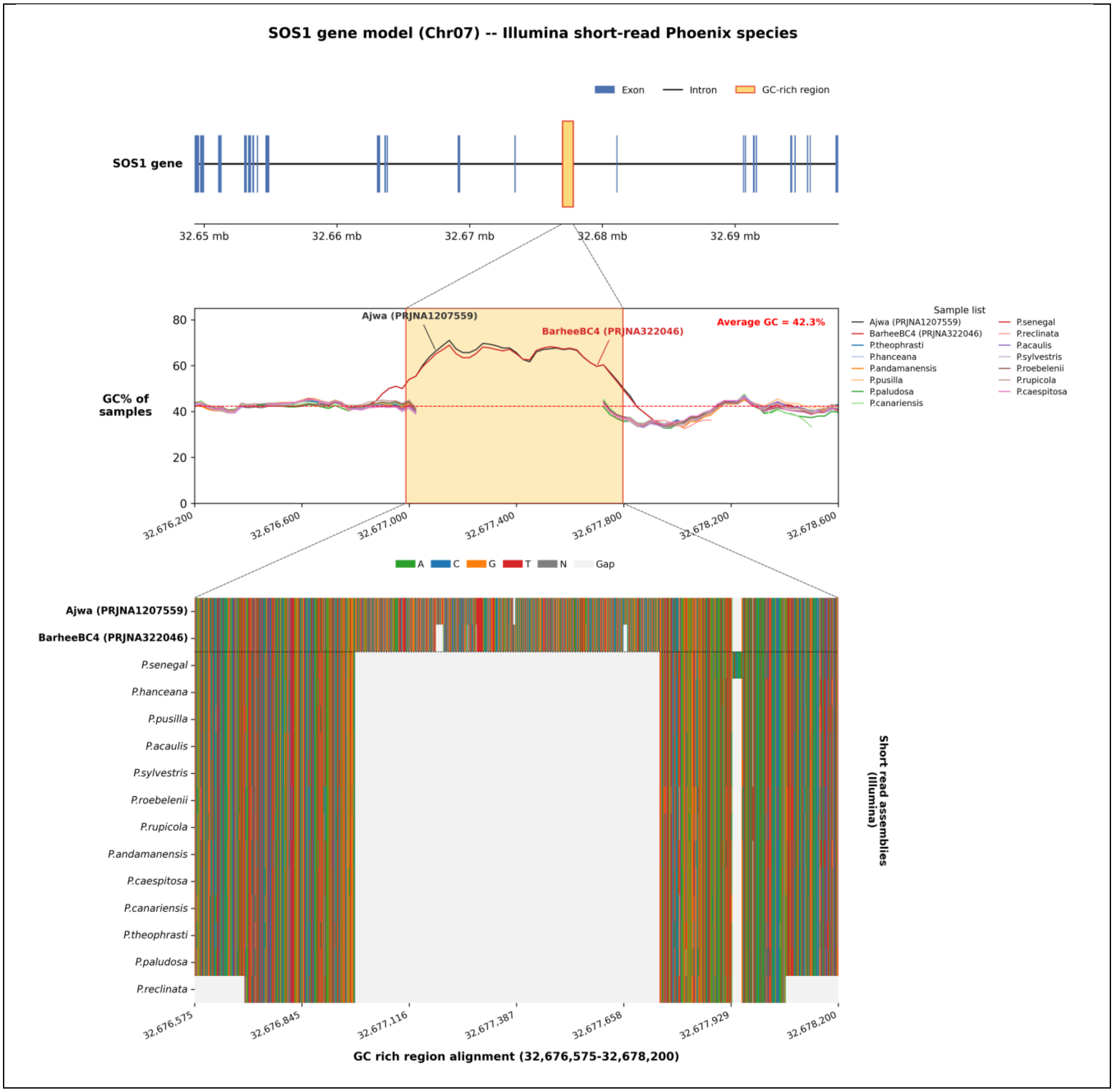

SOS1 gene model (Chr07) -- Illumina short-read Phoenix species
Exon
Intron
GC-rich region
SOS1 gene
32.65 mb
32.66 mb
32.67 mb
32.68 mb
32.69 mb
GC% of samples
Ajwa (PRJNA1207559)
BarheeBC4 (PRJNA322046)
Average GC = 42.3%
Sample list
Ajwa (PRJNA1207559)
BarheeBC4 (PRJNA322046)
P.theophrasti
P.hanceana
P.andamanensis
P.pusilla
P.paludosa
P.canariensis
P.senegal
P.reclinata
P.acaulis
P.sylvestris
P.roebelenii
P.rupicola
P.caespitosa
32,676,200
32,676,600
32,677,000
32,677,400
32,677,800
32,678,200
32,678,600
A
C
G
T
N
Gap
Ajwa (PRJNA1207559)
BarheeBC4 (PRJNA322046)
P.senegal
P.hanceana
P.pusilla
P.acaulis
P.sylvestris
P.roebelenii
P.rupicola
P.andamanensis
P.caespitosa
P.canariensis
P.theophrasti
P.paludosa
P.reclinata
Short read assemblies (Illumina)
32,676,575
32,676,845
32,677,116
32,677,387
32,677,658
32,677,929
32,678,200
GC rich region alignment (32,676,575-32,678,200)

Supplementary Figure S1. The pdSOS1 MITE insertion is specific to the date palm lineage. Top: gene model of pdSOS1 (Chr07), with exons (blue), introns (line) and the GC-rich MITE region (orange) within an intron. Middle: GC content across the region for the date palm reference assemblies Ajwa (PRJNA1207559) and BarheeBC4 (PRJNA322046) and the Phoenix species (genome-wide average GC = 42.3%, dashed line); the GC-rich MITE (shaded, Chr07:~32,677,000–32,677,800) forms a pronounced local GC peak in date palm. Bottom: nucleotide alignment of the GC-rich region (Chr07:32,676,575–32,678,200) across de novo short-read (Illumina) assemblies; coloured columns denote aligned bases (A, C, G, T; N in grey) and pale grey denotes gaps. The MITE insertion is present in both date palm assemblies (Ajwa, BarheeBC4) but absent, appearing as a gap, from all Phoenix species.

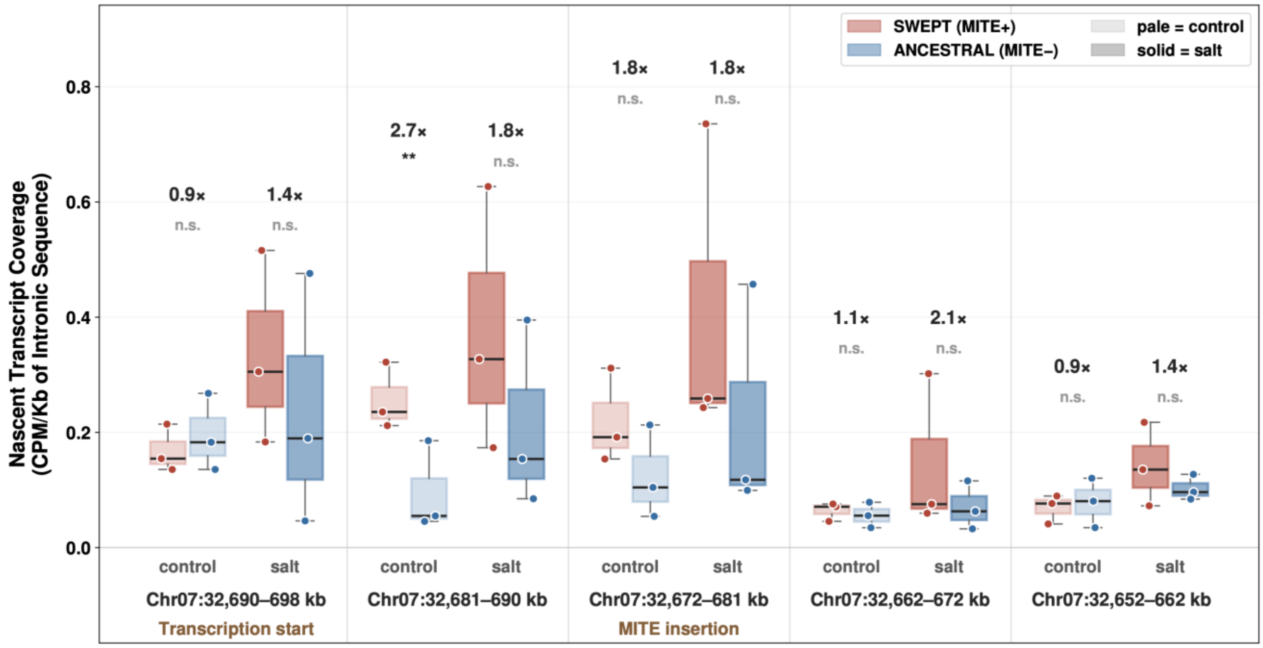


Supplementary Figure S2. pdSOS1 allele-specific gene expression analysis of swept and ancestral haplotypes in shoots. Gene expression data from date palms grown in control and high salinity conditions was analyzed for the cultivar Nabut Saif in shoots which contains both the ancestral (Mite-) and swept (Mite+) haplotypes. Allele-specific expression differences of nascent RNA (intronic) were observed with the swept haplotype demonstrating consistently higher levels of RNA in multiple 8kb windows of the gene. While differences are not always signficant they show a similar to trend to that observed in roots. Stars are significance based on paired T-test. Note that genome coordinates are in reverse order to match the gene transcription direction.

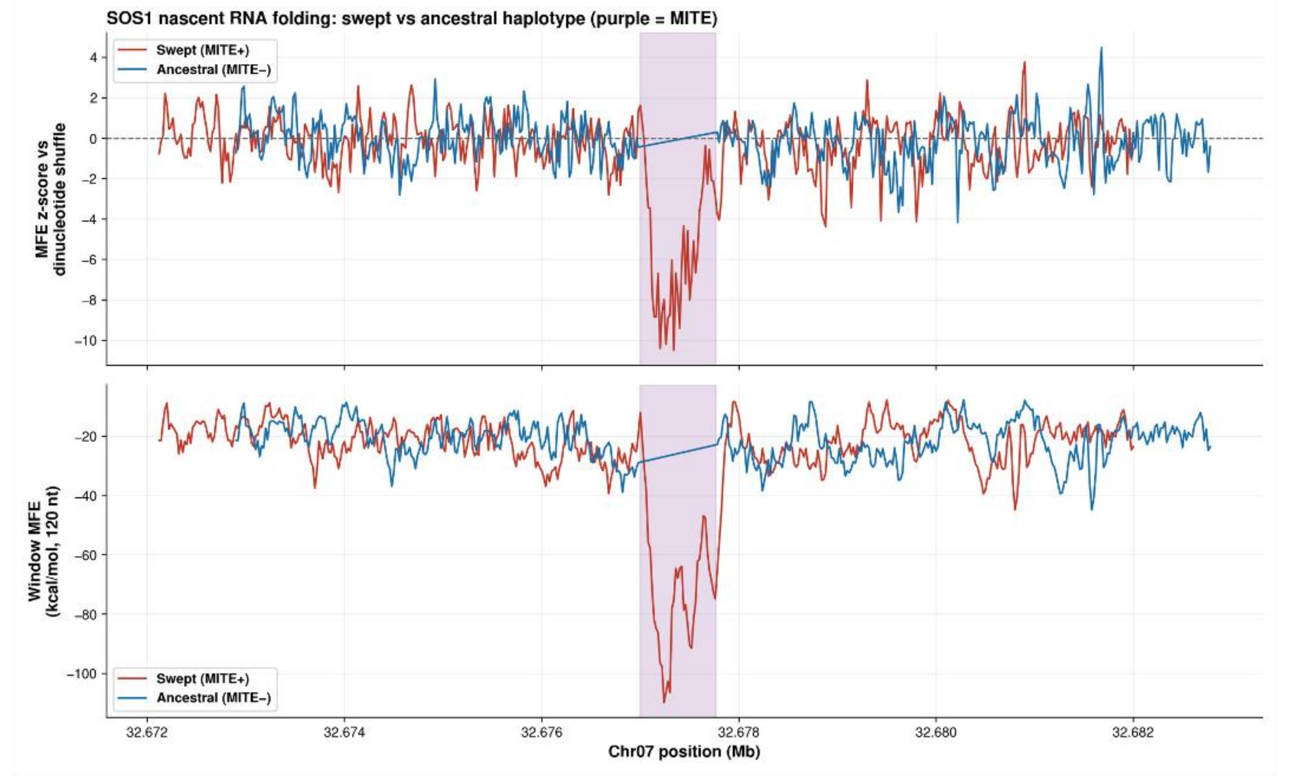


Supplementary Figure S3. RNA-folding analysis of the swept and non-Swept haplotype with MITE or shuffled insertion. ViennaFold 2.0 was used for minimum free energy analysis of the two haplotypes. The MITE inserted, swept haplotype or a dinucleotide shuffled version. The MITE insertion shows almost 10X the Minimum free energy in that region over the shuffled version suggesting a highly stable structure is possible in the RNA.

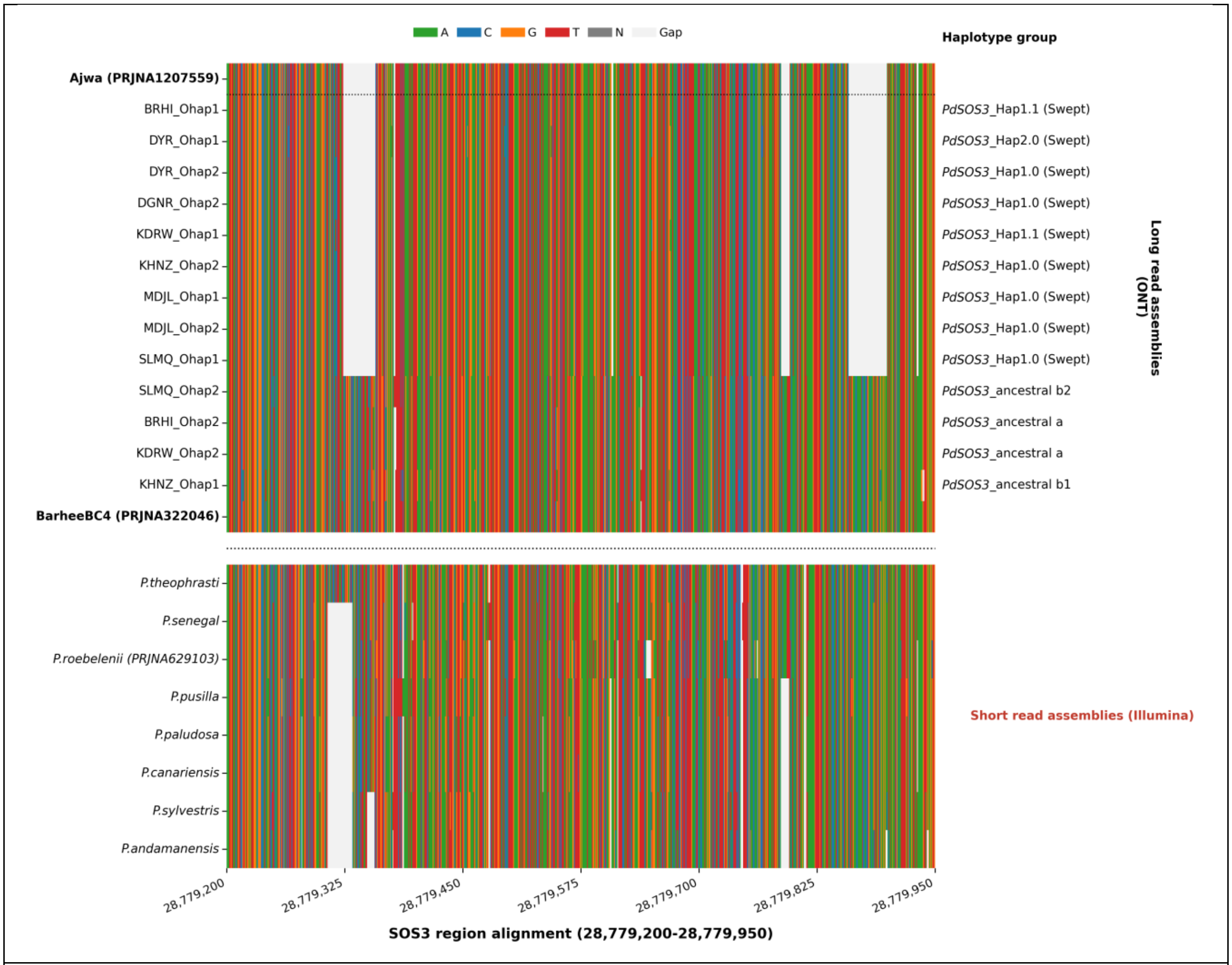


Supplementary Figure S4. Structural variation at the pdSOS3 promoter distinguishes swept from ancestral haplotypes. Nucleotide alignment of the *pdSOS3* promoter region (Chr09:28,779,200–28,779,950), spanning the 450-bp interspersed repeat (Dfam DR003745898.2; Chr09:28,779,351–28,779,801) and its two flanking deletions. **Top:** phased long-read (ONT) assemblies of date palm accessions (sample_Ohap1/2), labelled by haplotype group, swept (*pdSOS3*_hap1.0, hap1.1, hap2.0) and ancestral (anc-a, anc-b1, anc-b2), with the Ajwa (PRJNA1207559) and BarheeBC4 (PRJNA322046) references. **Bottom:** short-read (Illumina) assemblies of *Phoenix* species. Coloured columns denote aligned bases (A, C, G, T; N in grey); pale grey denotes gaps. The swept date palm haplotypes carry both deletions flanking the repeat, which are absent from the ancestral haplotypes. Across *Phoenix* species, the upstream deletion coincides with a target-site duplication of the repeat and varies among species, whereas the downstream deletion is specific to date palm.

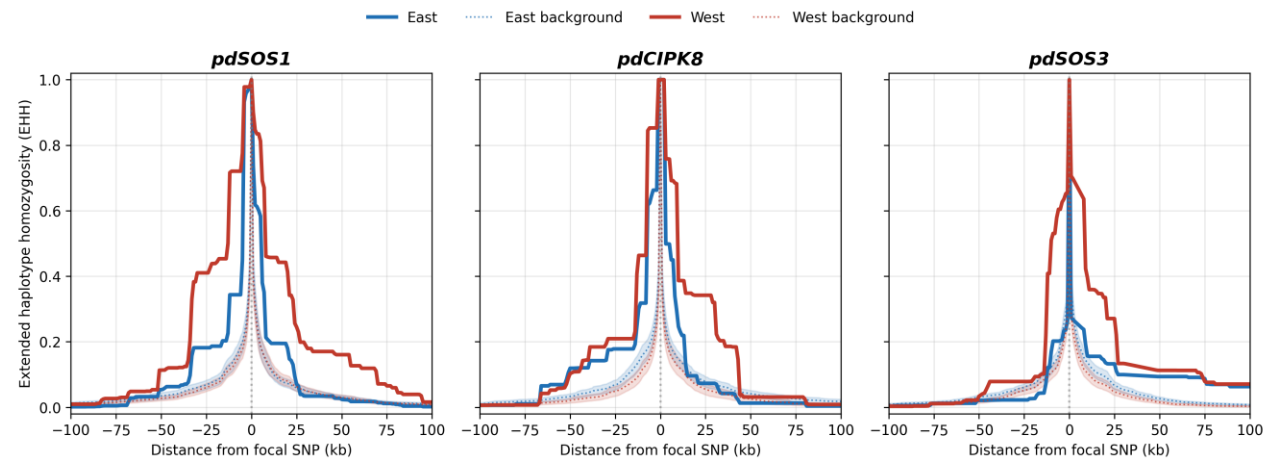


Supplementary Figure S5. Extended haplotype homozygosity (EHH) decay at the three swept gene regions, ±100 kb. EHH decay over a ±100 kb window around each focal SNP, *pdSOS1* (Chr07:32,679,064), *pdCIPK8* (Chr09:24,339,868) and *pdSOS3* (Chr09:28,779,415), in the Eastern (blue) and Western (red) date palm populations, with *P. theophrasti*-introgressed haplotypes excluded. Solid lines, focal EHH; dotted lines and shaded bands, the mean and 95% confidence interval of 100 allele-frequency–matched control SNPs across the chromosome. The wider window shows the full return of focal EHH to the background level beyond the swept interval.
